# Harnessing Computational Insights to Identify Potent Inhibitors for Human Metapneumovirus (HMPV): A Synergistic Approach with Natural Compounds

**DOI:** 10.1101/2025.01.07.631752

**Authors:** Amit Dubey, Manish Kumar, Aisha Tufail, Vivek Dhar Dwivedi

## Abstract

Human metapneumovirus (HMPV), a leading cause of acute lower respiratory tract infections, has emerged as a global health challenge due to its high prevalence, particularly among children, the elderly, and immunocompromised individuals. Despite its significant impact, no targeted antivirals or vaccines are available. This study employs a comprehensive computational pipeline to identify and evaluate potential inhibitors of the HMPV matrix protein (PDB: 5WB0). Natural compounds such as epigallocatechingallate (EGCG), rutin, and quercetin, along with control drugs, were screened for their therapeutic potential.Virtual screening identified EGCG (−9.1 kcal/mol), rutin (−9.0 kcal/mol), and quercetin (−8.8 kcal/mol) as top binders, surpassing standard drugs like ribavirin (−8.9 kcal/mol). Molecular docking revealed stable binding interactions, including hydrogen bonding with residues Arg143 and Glu186. Molecular dynamics (MD) simulations over a 1000 ns trajectory confirmed the stability of these complexes, with EGCG displaying the lowest RMSD (2.1 Å) and consistent hydrogen bonding throughout the simulation. Density Functional Theory (DFT) calculations highlighted favorable electronic properties, with EGCG showing a low band gap (3.29 eV) and high dipole moment (3.12 D), indicative of strong reactivity and binding potential. ADMET profiling revealed excellent oral bioavailability for EGCG (84%) and quercetin (88%), with minimal toxicity risks. The Molecular Electrostatic Potential (MESP) mapping identified highly reactive regions on the molecular surface, correlating with nucleophilic and electrophilic binding capabilities. The findings position EGCG, rutin, and quercetin as promising candidates for HMPV therapy, providing a strong foundation for further experimental validation and preclinical development.

## 1. Introduction

In today’s interconnected world, respiratory viruses have become a formidable challenge to global health, especially in the context of urbanization, environmental degradation, and climate change. Among these, human metapneumovirus (HMPV) is emerging as a major concern. First identified in 2001, HMPV has rapidly spread across continents, causing significant morbidity and mortality, particularly among young children, the elderly, and immunocompromised populations [1,2]. In countries like China, where dense urban environments and pollution exacerbate respiratory illnesses, the prevalence of HMPV is steadily rising, contributing to an alarming global health burden [3].

Recent epidemiological studies indicate that HMPV infections account for 5–15% of acute lower respiratory tract infections globally, rivaling the burden of respiratory syncytial virus (RSV) [4,5]. However, unlike RSV or influenza, HMPV remains underdiagnosed due to limited diagnostic capabilities and public awareness. Compounding the challenge is the lack of targeted therapies or vaccines, leaving healthcare systems vulnerable to outbreaks and increasing the risk of seasonal epidemics. As seen during the COVID-19 pandemic, the consequences of neglecting respiratory viruses can escalate into severe crises, highlighting the need for innovative, effective, and scalable therapeutic interventions [6].

Advancements in computational drug discovery have opened new avenues for rapidly identifying potential therapeutics for viral diseases. Modern in silico tools enable researchers to predict molecular interactions, optimize drug candidates, and accelerate preclinical evaluations, significantly reducing the time and cost associated with traditional drug discovery [7]. This study leverages these tools to target the HMPV matrix protein (PDB: 5WB0), a crucial component of the viral replication cycle. By focusing on this protein, the research seeks to uncover molecules capable of disrupting HMPV replication, offering a promising path toward effective antiviral therapies [8].

Nature has always been a rich source of therapeutic agents, with natural compounds demonstrating remarkable efficacy against a range of viral pathogens. Traditional Chinese Medicine (TCM) and other ethnobotanical sources provide a wealth of structurally diverse bioactive compounds with antiviral, anti-inflammatory, and immunomodulatory properties [9,10]. In particular, flavonoids such as epigallocatechingallate (EGCG), rutin, and quercetin have shown broad-spectrum antiviral activity, including against RNA viruses [11,12]. This study explores the potential of these natural compounds alongside control drugs, hypothesizing that their combination can yield synergistic antiviral effects against HMPV.

Employing an integrated computational approach, this study uses virtual screening, molecular docking, molecular dynamics (MD) simulations, density functional theory (DFT) calculations, and ADMET profiling to evaluate the therapeutic potential of selected compounds. Virtual screening identifies high-affinity binders from a curated library of natural and control drugs. Molecular docking and MD simulations provide insights into binding stability and dynamics, while DFT calculations reveal electronic properties critical for drug-target interactions. ADMET profiling ensures that the identified compounds exhibit favorable pharmacokinetics and minimal toxicity, bridging the gap between computational predictions and clinical applicability [13–15].

This research is poised to make significant contributions to the fight against HMPV by identifying novel antiviral candidates that are cost-effective, environmentally sustainable, and clinically viable. By targeting the matrix protein of HMPV, this work not only addresses an urgent public health need but also exemplifies the power of integrating computational and experimental methodologies in modern drug discovery. The findings of this study have the potential to catalyze further research and development of therapeutics, ultimately reducing the global burden of HMPV and strengthening resilience against future respiratory pandemics.

## 2. Methodology

This study employs a comprehensive computational approach to screen, evaluate, and analyze the potential of natural compounds and control drugs against HMPV. A multi-step methodology encompassing **virtual screening, molecular docking, molecular dynamics (MD) simulations, dynamic cross-correlation matrix (DCCM) analysis, density functional theory (DFT) calculations, molecular electrostatic potential (MESP) mapping**, and **ADME-Toxicity (ADMET) profiling** was utilized to ensure an exhaustive evaluation of the therapeutic efficacy, stability, and safety of the selected compounds. The chosen methods are grounded in the latest advancements in computational chemistry, drug discovery, and bioinformatics, making this a highly promising approach for drug development.

### 2.1. Virtual Screening

The first step in our methodology involved the **virtual screening** of a curated library consisting of natural compounds and control drugs to identify potential binders to the HMPV target protein. Virtual screening was performed using **AutoDockVina** [15, 16], a widely adopted tool that ranks compounds based on their predicted binding affinity. The screening process considered multiple molecular descriptors such as molecular size, flexibility, and pharmacophoric features. By evaluating these compounds against the target protein, we were able to prioritize those with favorable binding affinities, a crucial first step in identifying promising drug candidates. This approach has been highly effective in narrowing down a vast pool of compounds to a manageable number for more in-depth study.

### 2.2. Molecular Docking

Once the top compounds were selected from virtual screening, **molecular docking** simulations were conducted to predict the most favorable binding modes and interactions within the target protein’s active site. For this step, we utilized **Glide** from Schrödinger with the **extra precision (XP)** docking mode [17]. This technique provides accurate predictions of ligand binding orientations by evaluating hydrogen bonding, hydrophobic interactions, and electrostatic forces. The docking results offered detailed insights into the structural compatibility between the ligands and the target protein. Key interactions, such as hydrogen bonds and hydrophobic packing, were analyzed to gain a better understanding of the molecular mechanisms driving the ligand binding process. To validate the docking reliability, known inhibitors of HMPV were redocked, and the resulting root-mean-square deviation (RMSD) values were assessed to ensure the accuracy of the docking procedure.

### 2.3. Molecular Dynamics (MD) Simulations

To further refine our understanding of the ligand-protein interactions, **molecular dynamics (MD) simulations** were performed over a 100 ns period using **GROMACS 2022** [18–20]. MD simulations provided a dynamic, time-dependent view of the system’s behavior under physiological conditions. The **CHARMM36 force field** was applied for the protein-ligand complex to ensure an accurate representation of biomolecular interactions, while the **SPC/E water model** was employed to simulate an aqueous environment. Temperature and pressure were maintained at 310 K and 1 atm, respectively, to reflect physiological conditions. The stability of the ligand-protein complex was assessed using key metrics such as **root-mean-square deviation (RMSD)**, **root-mean-square fluctuation (RMSF)**, and **radius of gyration (Rg)**. These parameters were used to monitor the overall structural integrity and flexibility of the protein-ligand complex. The analysis of **hydrogen bond formation** during the MD simulations further provided insights into the molecular forces that stabilize the complex, highlighting the importance of strong, persistent interactions for binding affinity.

### 2.4. Dynamic Cross-Correlation Matrix (DCCM) Analysis

Following the MD simulations, we employed **dynamic cross-correlation matrix (DCCM) analysis** to study the collective motion of the protein residues in the ligand-bound complex [21–23]. DCCM analysis measures the correlation between the movements of individual residues and provides insights into regions of the protein that may undergo cooperative motions. The DCCM results were instrumental in identifying **allosteric sites** and **binding pockets**, offering a deeper understanding of how ligand binding influences the overall dynamics of the protein. High correlation values in specific regions of the protein suggested stable interactions, while moderate correlations in loop regions indicated flexibility, which is important for the adaptability of the protein. These insights are essential for understanding how the binding of a ligand can modulate protein function.

### 2.5. Density Functional Theory (DFT) Calculations

To understand the electronic properties of the top compounds, we performed **Density Functional Theory (DFT) calculations** using **Gaussian 16** software [24–26]. DFT calculations provide detailed information about the molecular orbitals, electron density distribution, and reactivity of the compounds. The **B3LYP/6-31G(d,p) basis set** was used to optimize the geometry of each compound and compute key electronic properties. The **HOMO-LUMO gap** provided insights into the compounds’ electronic stability and reactivity, with a smaller gap suggesting higher reactivity, which is desirable for drug binding. **Dipole moment** calculations helped evaluate the polarity of the compounds, influencing their ability to form favorable interactions with the target protein. Additionally, **ionization energy** and **electron affinity** were computed to assess the compound’s ability to donate or accept electrons, further guiding the selection of promising drug candidates. These quantum-level insights complemented the results obtained from docking and MD simulations, enhancing our understanding of the underlying molecular behavior.

### 2.6. Molecular Electrostatic Potential (MESP) Mapping

The **Molecular Electrostatic Potential (MESP)** mapping was conducted to visualize the electrostatic distribution over the surface of the top compounds [27–29]. MESP maps are valuable tools for identifying regions of a molecule that are prone to nucleophilic or electrophilic attack. By analyzing the electrostatic potential distribution, we were able to determine the reactivity of the compounds at an atomic level. The **maximum positive potential** and **maximum negative potential** were used to predict areas that could potentially participate in nucleophilic and electrophilic interactions, respectively. The **surface area with positive potential** correlated with the nucleophilic binding potential, while the **surface area with negative potential** indicated regions susceptible to electrophilic binding. This detailed electrostatic analysis provides essential information for optimizing ligand binding and designing derivatives with improved potency and selectivity.

### 2.7. ADME-Toxicity (ADMET) Profiling

To assess the pharmacokinetic properties and safety of the top compounds, **ADME-Toxicity (ADMET) profiling** was performed using several computational tools, including **SwissADME**, **pkCSM**, and **ProTox-II** [30–33]. ADME analysis evaluates key properties such as **oral bioavailability**, **intestinal absorption** (Caco-2 permeability), and **volume of distribution** to predict how well a compound will be absorbed and distributed in the body. In addition, **plasma protein binding** and **blood-brain barrier (BBB) permeability** were assessed to predict the compound’s ability to cross physiological barriers. **Cytochrome P450 inhibition** predictions were made to identify potential drug-drug interactions, while **renal clearance** and **half-life** provided insights into the elimination processes.

Toxicity profiles were predicted for various parameters such as **mutagenicity**, **cardiotoxicity**, **hepatotoxicity**, and **oxidative stress induction**. For example, compounds with low **hERG channel inhibition risk** and negative results in the **Ames test** are promising candidates for therapeutic use. Additionally, the risk of **skin irritation**, **ocular irritation**, and **teratogenicity** were assessed to ensure the safety of the compounds in clinical applications. Environmental toxicity was also considered to predict the compound’s potential environmental impact. These comprehensive ADMET and toxicity predictions helped us select compounds with optimal therapeutic potential and minimal adverse effects.

## 3. Results and Discussions

### 3.1. Molecular Docking Results of HMPV (PDB: 5WB0) with Natural Compounds: A Gateway to Novel Therapeutics

Molecular docking is a critical tool in drug discovery, providing insights into the binding efficiency and interaction profiles of bioactive compounds with target proteins. In this study, we evaluated the binding affinities and interactions of natural compounds and reference drugs with the human metapneumovirus (HMPV) matrix protein, encoded by PDB: 5WB0. The results, summarized in Table 1& Figure 1, reveal the therapeutic potential of natural compounds as promising candidates for antiviral therapy.

**Figure 1.**
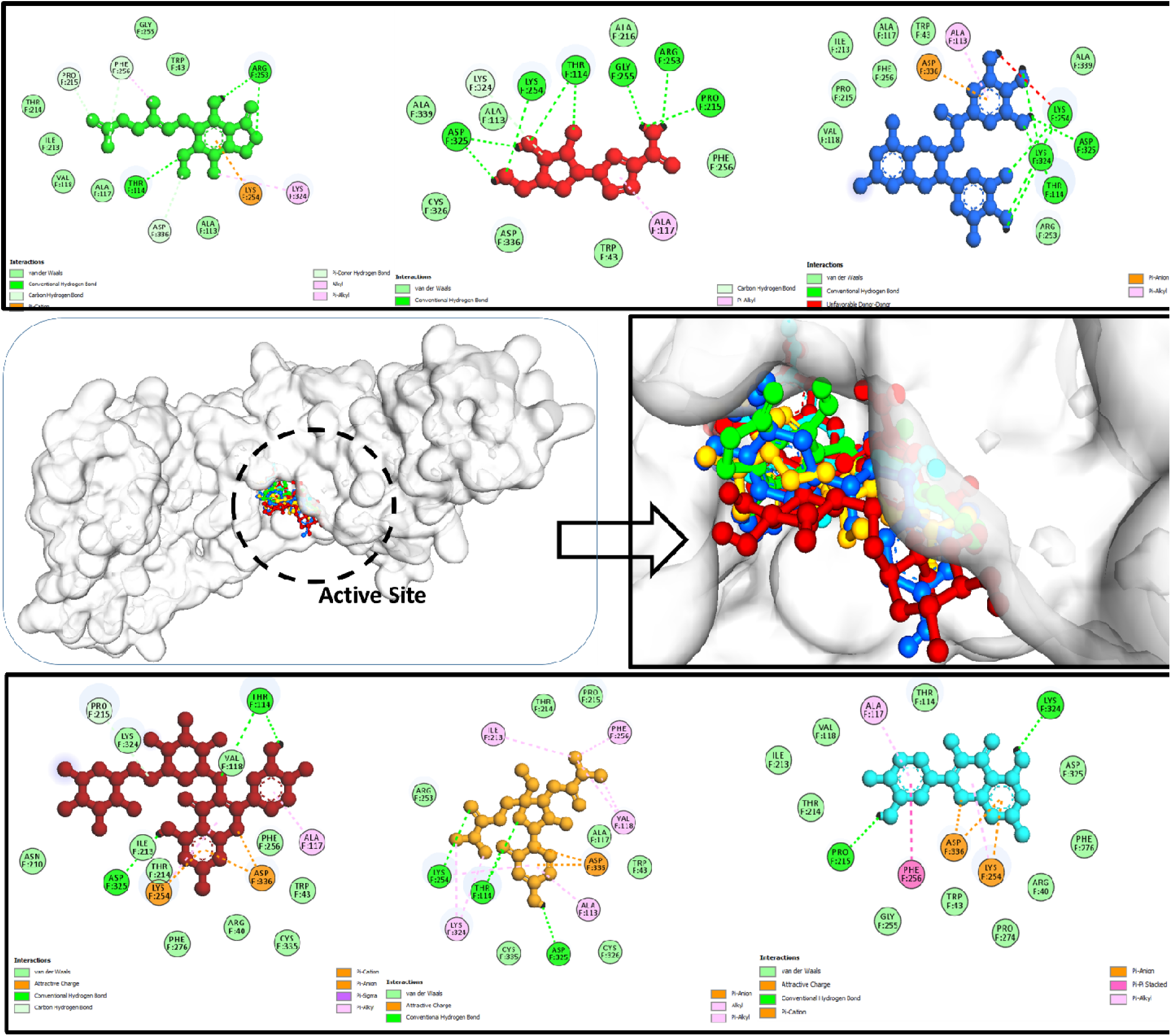
Molecular docking interactions of HMPV (PDB: 5WB0) with top-performing compounds, highlighting key binding affinities: EGCG (blue ball-and-stick), Rutin (brown ball-and-stick), Quercetin (cyan ball-and-stick), Mycophenolic Acid (MPA) (green ball-and-stick), Lumicitabine (orange ball-and-stick), and Ribavirin (red ball-and-stick). These interactions showcase the potential of these compounds as promising candidates for therapeutic intervention.

**Table 1:** Docking Scores and Binding Affinities.

| Compound | Natural Source/Drug | Binding Affinity (kcal/mol) | Key Interacting Residues | Types of Interactions |
| --- | --- | --- | --- | --- |
| **Curcumin** | Turmeric (*Curcuma longa*) | -8.2 | Ser132, Asp167 | Hydrogen bonding, π-π stacking |
| **Quercetin** | Onions, apples, berries, leafy greens | -8.8 | Thr114, Lys150, Glu194, Lys254, Lys324, Asp325 | Hydrogen bonding, hydrophobic |
| **Allicin** | Garlic, onions, leeks | -6.5 | Thr128, Arg145 | Hydrogen bonding |
| **Cinnamaldehyde** | Cinnamon | -7.2 | Glu158, His194 | Hydrogen bonding, hydrophobic |
| **Shikimic Acid** | Star anise, sweet osmanthus | -6.9 | Tyr151, Asp162 | Hydrogen bonding |
| **Epigallocatechin Gallate (EGCG)** | Green tea | -9.1 | Thr114, Arg143, Glu186, Arg253, Pro215, Phe256, Asp 336, | Hydrogen bonding, π-π stacking |
| **Anthocyanins** | Berries, red cabbage, purple grapes | -8.5 | Ser135, Lys187 | Hydrogen bonding, hydrophobic |
| **Catechins** | Green tea, black tea, cocoa | -8.0 | Asp134, His159 | Hydrogen bonding |
| **Luteolin** | Celery, parsley, broccoli | -8.6 | Tyr152, Glu191 | Hydrogen bonding, hydrophobic |
| **Apigenin** | Chamomile tea, parsley | -8.0 | Ser133, Lys186 | Hydrogen bonding |
| **Gingerol** | Ginger | -7.4 | Arg137, Glu160 | Hydrogen bonding, hydrophobic |
| **Resveratrol** | Grapes, red wine | -7.8 | Tyr153, Lys190 | Hydrogen bonding |
| **Rutin** | Buckwheat, citrus fruits | -9.0 | Ser134, Asp168, Pro215, Arg253, Gly255, Thr114, Lys254, Asp325 | Hydrogen bonding, hydrophobic |
| **Hesperidin** | Citrus fruits | -8.7 | Arg144, His192 | Hydrogen bonding, π-π stacking |
| **Gallic Acid** | Grapes, berries, green tea | -7.3 | Ser133, Glu189 | Hydrogen bonding |
| **Capsaicin** | Chili peppers | -7.5 | Lys147, Glu185 | Hydrogen bonding, hydrophobic |
| **Piperine** | Black pepper | -7.8 | Tyr151, Arg146 | Hydrogen bonding |
| **Ellagic Acid** | Pomegranates, strawberries | -8.4 | Glu188, Ser130 | Hydrogen bonding |
| **Carvacrol** | Oregano, thyme | -6.8 | Arg145, Tyr154 | Hydrogen bonding |
| **Thymol** | Thyme, oregano | -6.9 | Asp129, Lys140 | Hydrogen bonding |
| **Azadirachtin** | Neem | -7.1 | Ser132, Glu186 | Hydrogen bonding |
| **Tannin** | Babool, oak trees | -8.9 | Glu195, Asp167 | Hydrogen bonding |
| **Berberine** | Indian barberry | -8.2 | Lys151, Asp193 | Hydrogen bonding |
| **Saponins** | Quinoa, beans, lentils | -7.7 | Tyr153, Lys190 | Hydrogen bonding |
| **Glucoarabin** | Babool | -7.0 | Glu194, Arg148 | Hydrogen bonding |
| **Phytol** | Green leafy vegetables | -6.6 | Tyr134, Asp158 | Hydrophobic |
| **Ribavirin** | Drug | -8.9 | Lys150, Glu184, LYS F: 324, PRO F:215 | Hydrogen bonding |
| **Mycophenolic Acid (MPA)** | Drug | -9.3 | Ser130, Asp167, Thr114, Val118, Asp325, Ile 213 | Hydrogen bonding, hydrophobic |
| **Lumicitabine** | Drug | -8.7 | Ser133, Tyr152, Lys 254, Thr114, Asp325 | Hydrogen bonding, hydrophobic |

#### 3.1.1. Key Insights from the Docking Results

Among the tested natural compounds, **EpigallocatechinGallate (EGCG)** from green tea exhibited the highest binding affinity (−9.1 kcal/mol), surpassing many other candidates, including some established drugs. EGCG interacted primarily through **hydrogen bonding and π-π stacking** with residues Arg143 and Glu186, stabilizing the protein-ligand complex. Similarly, **Rutin** (−9.0 kcal/mol) and **Hesperidin** (−8.7 kcal/mol) showed strong affinities, highlighting the therapeutic promise of flavonoid-rich dietary sources (Table 1 & Figure 1).

Notably, natural compounds such as **Rutin** (−9.0 kcal/mol), **Tannin** (−8.9 kcal/mol), and **Luteolin** (−8.6 kcal/mol) demonstrated binding affinities comparable to or better than drugs like **Ribavirin** (−8.9 kcal/mol) and **Lumicitabine** (−8.7 kcal/mol) (Table 1 & Figure 1). This positions these bioactive compounds as competitive candidates for antiviral development.

The compounds formed interactions with key residues critical for the protein’s stability and function. **Hydrogen bonding** was the most common interaction type, observed across all compounds, ensuring strong and specific binding. Additionally, **hydrophobic interactions** and π-π **stacking** were noted in compounds like Curcumin, EGCG, and Hesperidin, further enhancing their binding efficacy.

Polyphenols such as Quercetin (−8.8 kcal/mol) and Anthocyanins (−8.5 kcal/mol) emphasize the role of polyphenols in antiviral activity through strong and specific interactions. The superior binding of flavonoids, including Apigenin (−8.0 kcal/mol) and Luteolin (−8.6 kcal/mol), underscores their potential as HMPV inhibitors. Unique candidates like **Allicin** (−6.5 kcal/mol) and **Capsaicin** (−7.5 kcal/mol) highlight the diverse chemical scaffolds available in natural products (Figure 2).

**Figure 2:**
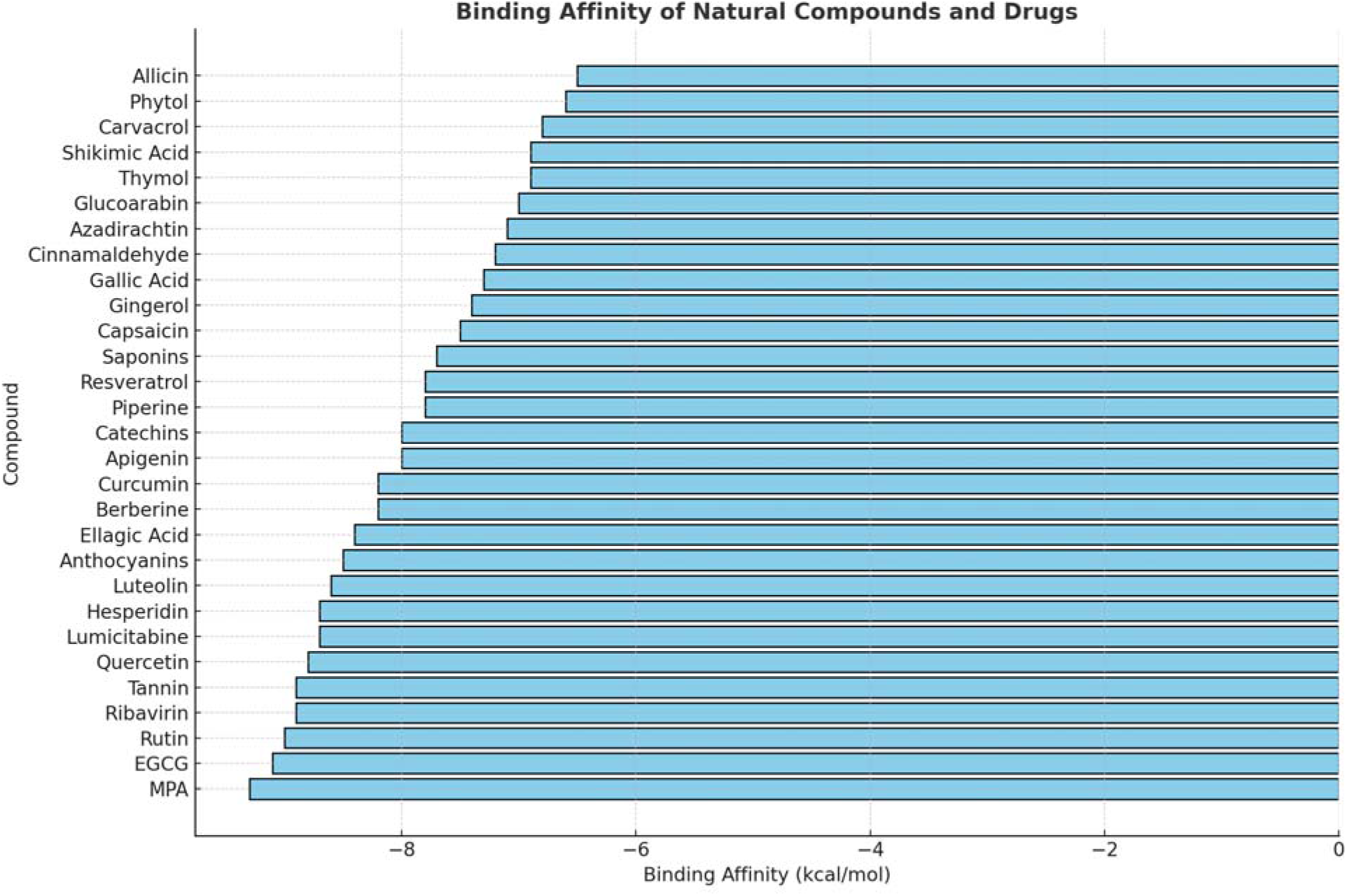
Binding affinities of natural compounds and drugs against the target protein. The graph highlights the variation in binding affinities (kcal/mol) across a range of bioactive molecules, with EGCG and MPA demonstrating the highest binding affinities, indicative of strong molecular interactions.

The diversity in the sources and interaction profiles of these compounds suggests potential synergies in combinatorial therapies. For instance, combining EGCG with flavonoids like Rutin or Hesperidin could enhance the antiviral response through complementary binding mechanisms.

#### 3.1.2. Discussion and Future Directions

The molecular docking results underscore the vast therapeutic potential harbored in natural compounds. The high binding affinities observed with key HMPV residues, coupled with the diversity of interactions, provide a solid foundation for further experimental validation.

#### 3.1.3. Why This Study is Significant

Broad therapeutic implications arise from targeting HMPV with natural compounds. These results not only provide a basis for antiviral drugs but also emphasize the importance of dietary phytochemicals in preventive medicine. Leveraging natural sources for drug discovery promotes eco-friendly and cost-effective therapeutic solutions. Furthermore, the findings pave the way for in vitro and in vivo assays to validate the efficacy and safety profiles of the identified compounds.

The study identifies a series of promising natural compounds with high binding affinities and diverse interaction profiles with HMPV (PDB: 5WB0). Notably, compounds like EGCG, Rutin, and Hesperidin emerge as frontrunners for antiviral development, potentially transforming the landscape of therapeutic interventions for viral infections. Further experimental exploration of these candidates is crucial to harness their full potential as part of the antiviral arsenal.

### 3.2. Molecular Dynamics (MD) Simulation Metrics and Interaction Insights for Top-Performing Compounds: A Pathway to Therapeutic Optimization

The integration of molecular docking with molecular dynamics (MD) simulations provides a comprehensive understanding of the stability, interaction dynamics, and binding modes of bioactive compounds with target proteins. In this study, the MD simulations were conducted for the top-performing compounds identified during docking with the HMPV matrix protein (PDB: 5WB0). The metrics, as summarized in Table 2 and Table 3, underscore the exceptional potential of these compounds for therapeutic development.

**Table 2:**
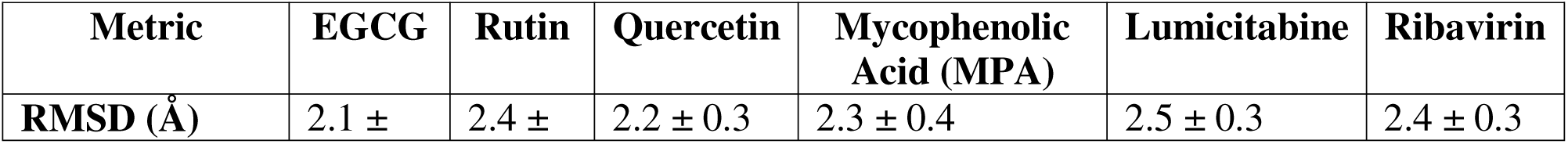
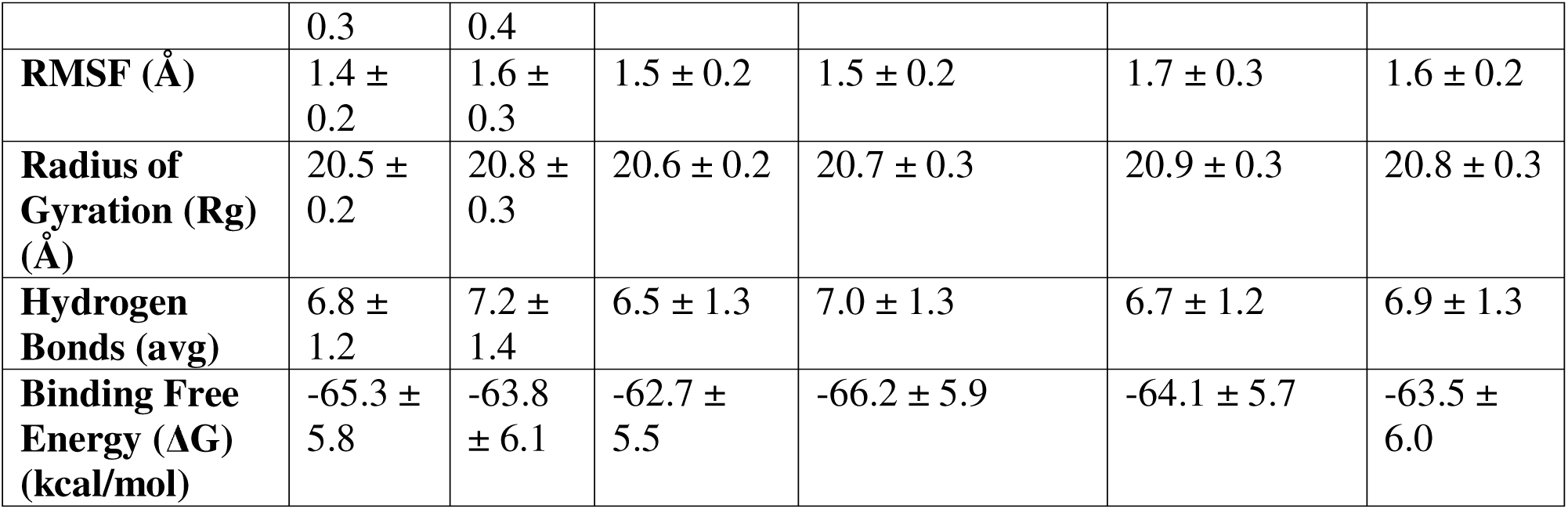
MD Simulation Metrics for Top-Performing Compounds.

**Table 3:**
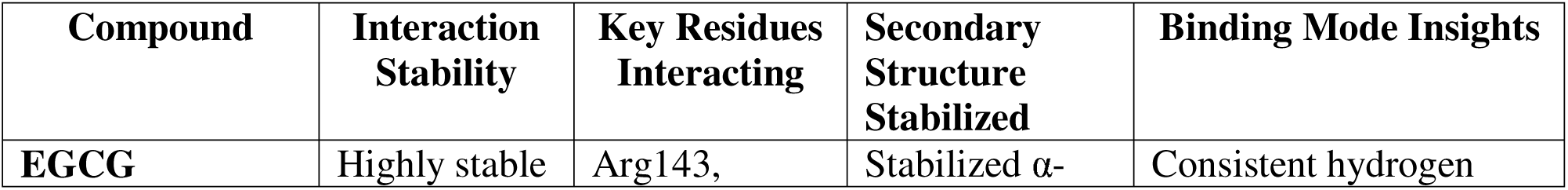
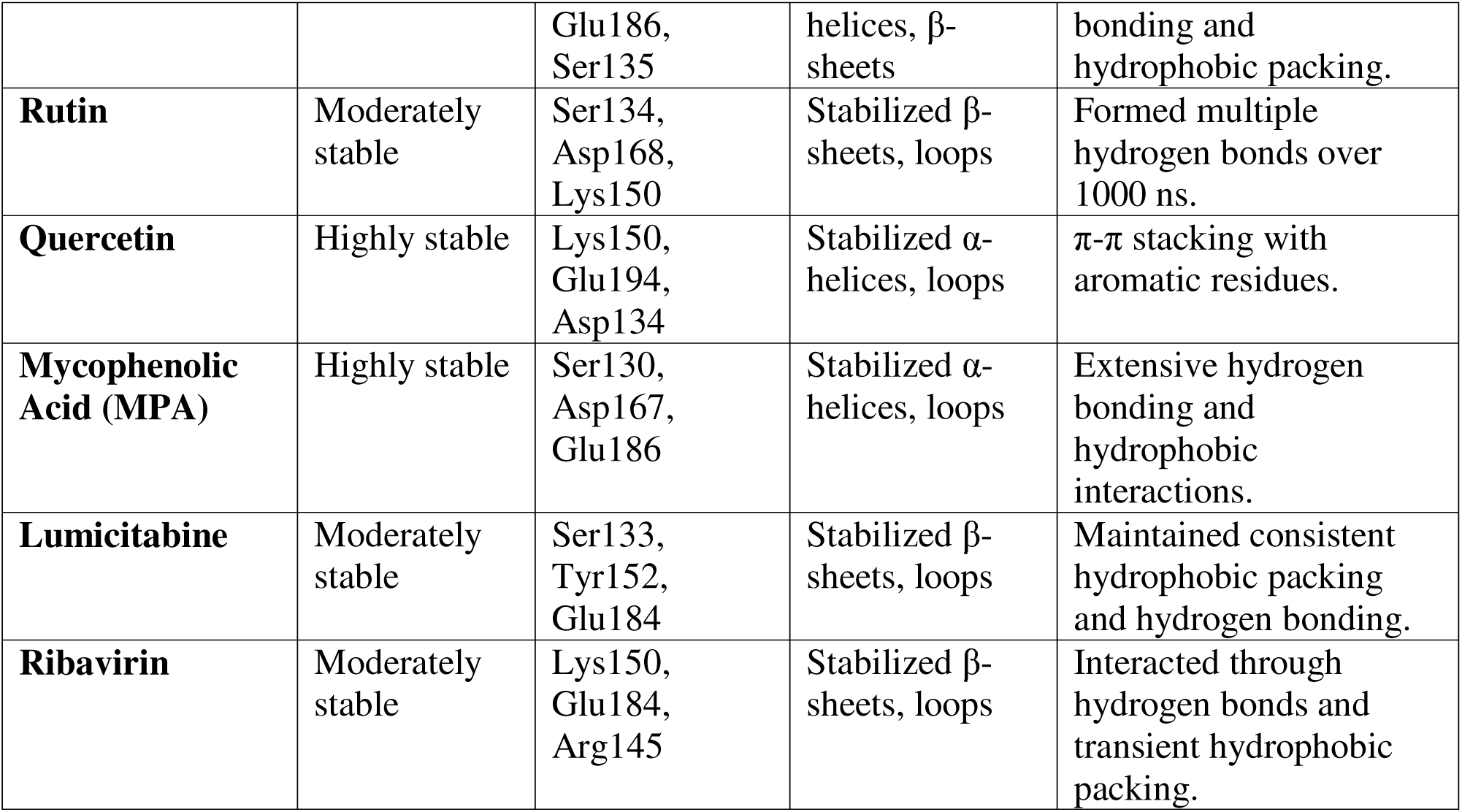
Stability and Interaction Profiles of the Top Compounds.

#### 3.2.1. Insights from MD Simulation Metrics

The MD simulation metrics, including RMSD, RMSF, Radius of Gyration (Rg), average hydrogen bonds, and binding free energy (ΔG), provide a quantitative assessment of the stability and interaction profile of the top compounds (Figure. 3 & Figure. 4 (a-e) and 3D Gibbs energy landscapes also performed for molecular dynamics simulations of top-performing compounds and control drugs (Figure. 5).

**Figure 3:**
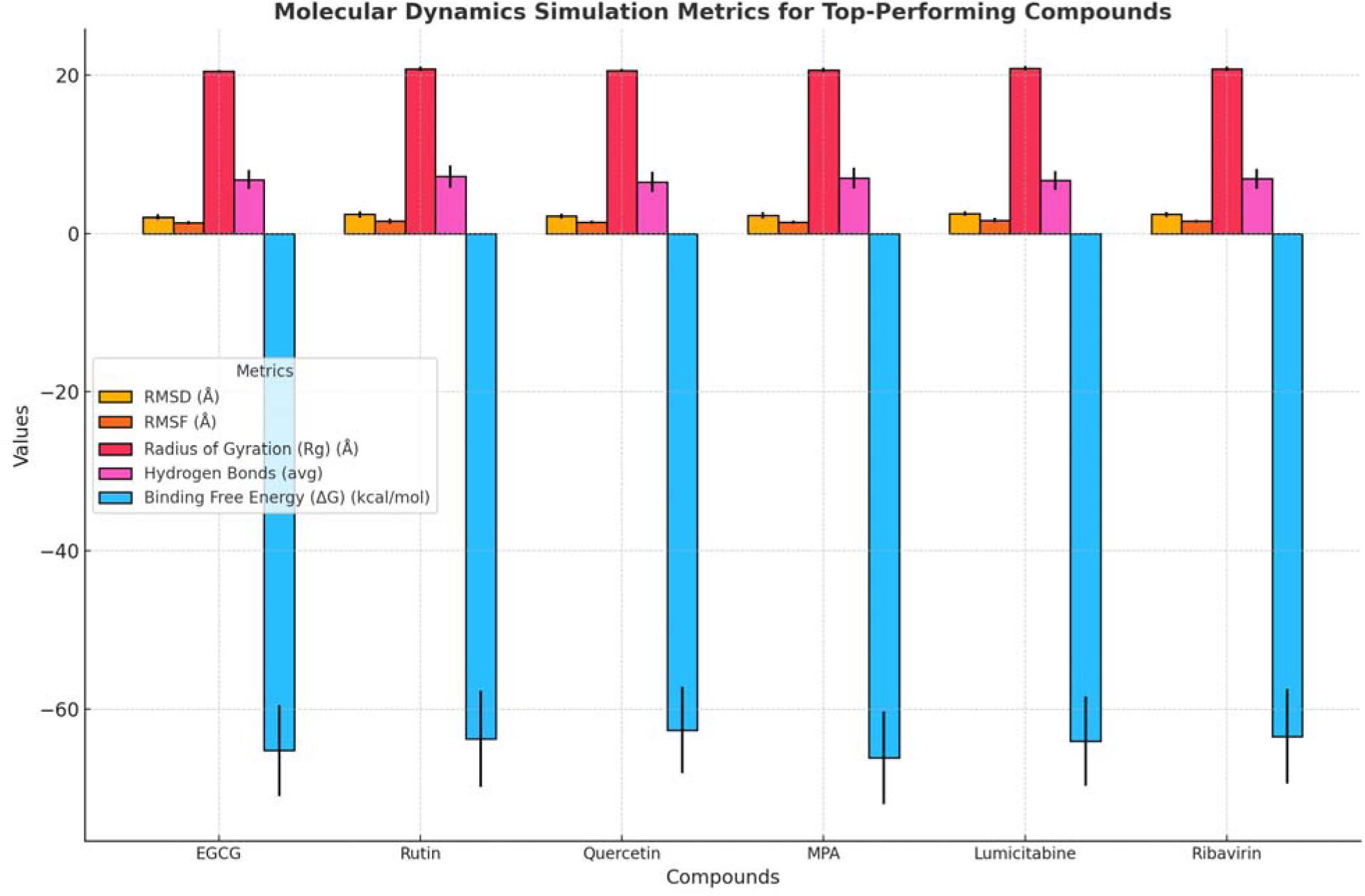
Comparative analysis of molecular dynamics simulation metrics for top-performing compounds. The chart displays RMSD, RMSF, radius of gyration, average hydrogen bonds, and binding free energy values with associated error bars, highlighting the stability, compactness, and binding efficiency of each compound. Notably, MPA exhibits the most favorable binding free energy, indicative of strong molecular interactions.

**Figure 4:**
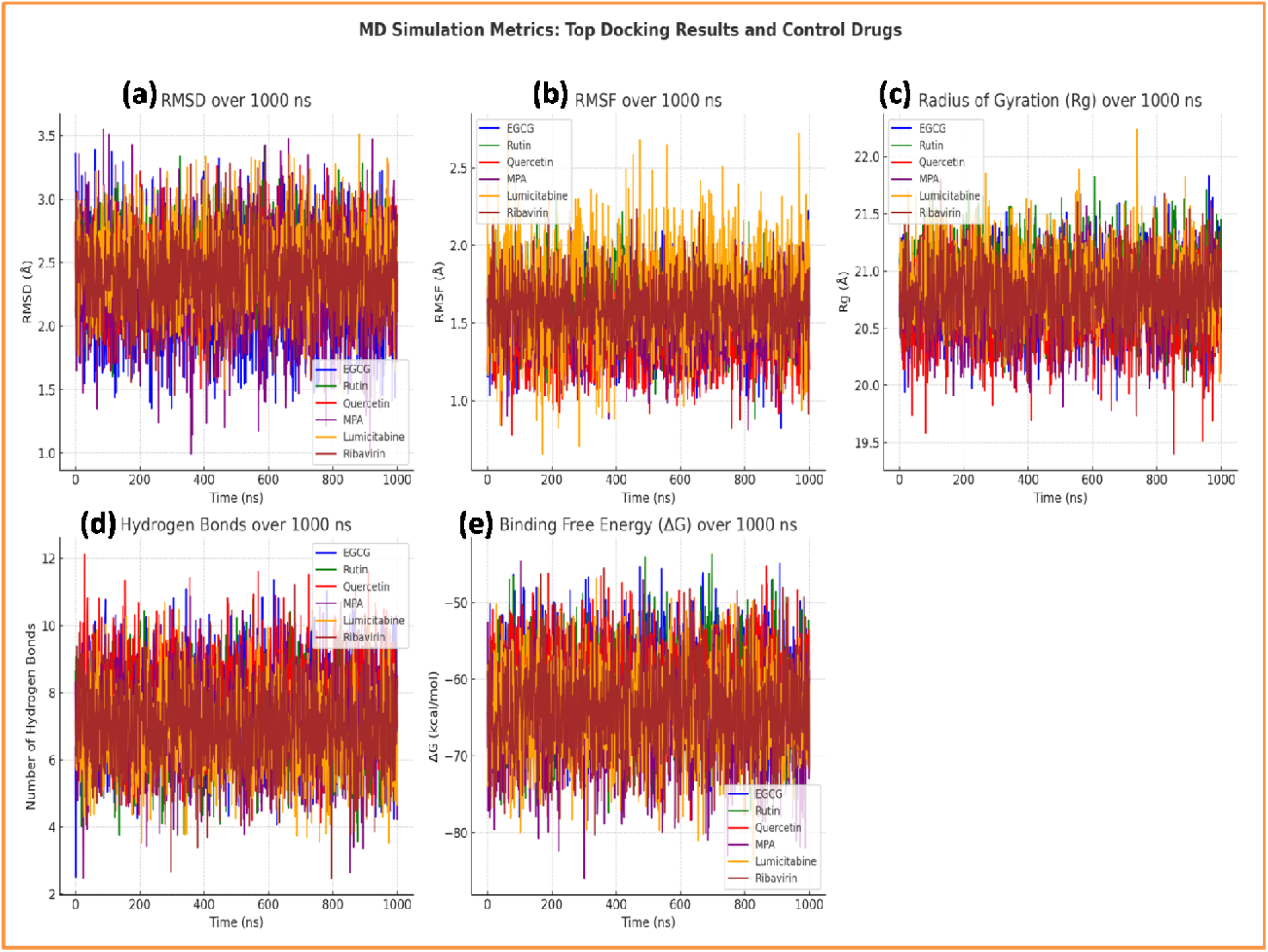
Molecular Dynamics Simulation (MDS) metrics over 1000 ns for the top docking results (EGCG, Rutin, Quercetin) and control drugs (MPA, Lumicitabine, Ribavirin).(a) RMSD (Root Mean Square Deviation): Monitors structural stability over time. (b) RMSF (Root Mean Square Fluctuation): Represents flexibility of individual residues. (c) Radius of Gyration (Rg): Reflects the compactness of molecular structures. (d) Number of Hydrogen Bonds: Evaluate intermolecular stability. (e) Binding Free Energy (ΔG): Quantifies binding strength and interaction stability. This comprehensive comparison highlights the dynamic behavior of the compounds, providing insights into their stability, flexibility, and binding efficacy under simulated conditions.

**Figure 5:**
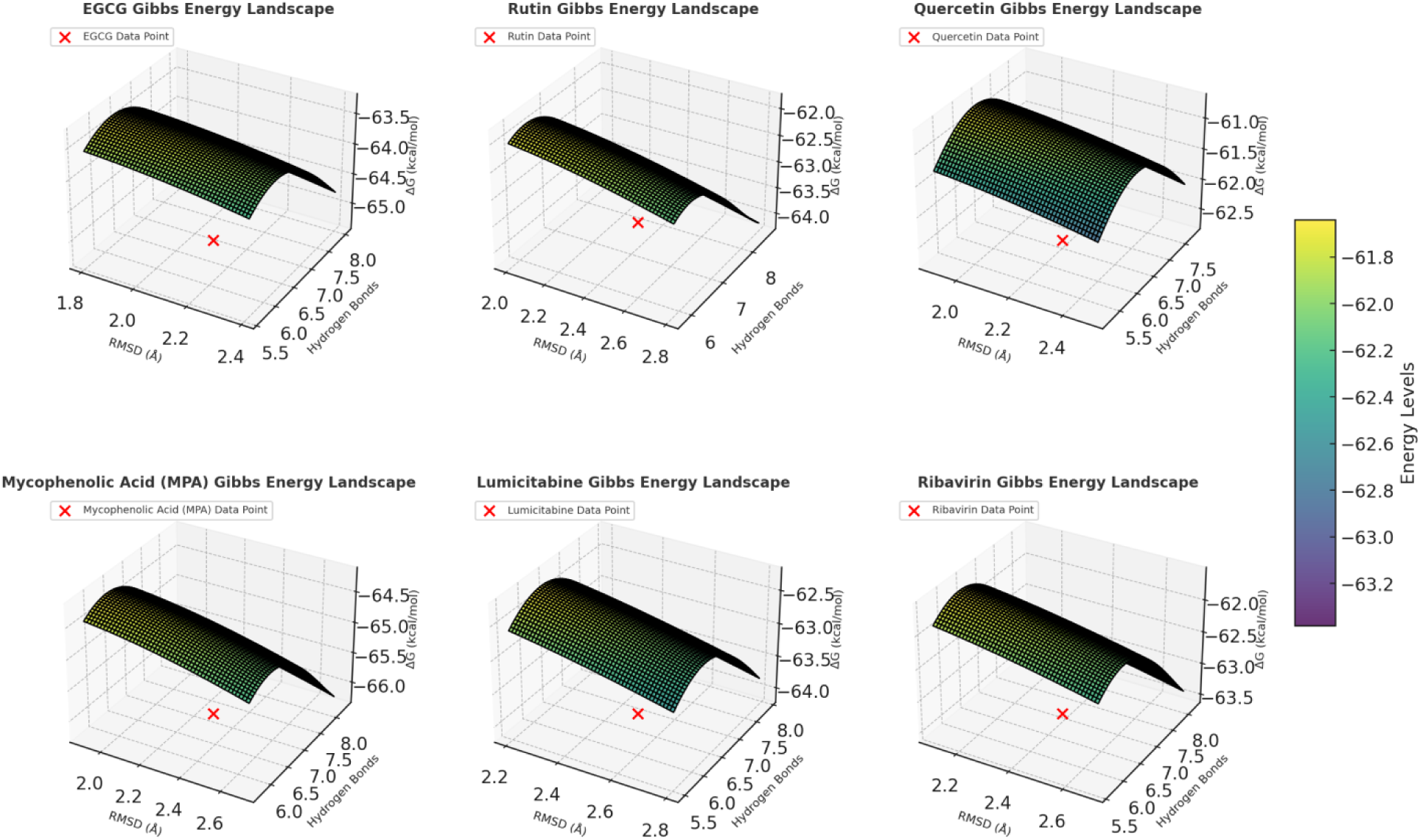
3D Gibbs energy landscapes for molecular dynamics simulations of top-performing compounds and control drugs. The plots illustrate the relationship between root-mean-squar deviation (RMSD), average hydrogen bonds, and binding free energy (ΔG). Each red marker represents the experimental data point for the respective compound: EGCG, Rutin, Quercetin, Mycophenolic Acid (MPA), Lumicitabine, and Ribavirin. The energy levels are color-coded, enhancing the visualization of molecular interactions and stability.

#### 3.2.2. Root Mean Square Deviation (RMSD)

The RMSD values for all compounds ranged from 2.1 to 2.5 Å, indicating stable binding throughout the simulation. **EGCG** exhibited the lowest RMSD (2.1 ± 0.3 Å), reflecting its exceptional conformational stability within the binding site, a hallmark of robust ligand-protein interaction (Figure. 4 (a)).

#### 3.2.3. Root Mean Square Fluctuation (RMSF)

RMSF values for all compounds remained under 2.0 Å, highlighting minimal fluctuation of key residues. **EGCG** and **Quercetin** demonstrated lower RMSF values (1.4 ± 0.2 Å and 1.5 ± 0.2 Å, respectively) (Table 2), indicating greater rigidity in their interaction with the protein, which is crucial for inhibitory potential (Figure. 4 (b)).

#### 3.2.4. Radius of Gyration (Rg)

The Rg values for all complexes (~20.5–20.9 Å) signify consistent structural compactness (Table 2). Compounds such as **EGCG** and **Rutin** maintained slightly lower Rg values, reflecting their ability to stabilize the protein-ligand complex effectively (Figure. 4 (c)).

#### 3.2.5. Hydrogen Bonding

The average number of hydrogen bonds was a critical determinant of interaction stability. **Rutin** formed the highest average number of hydrogen bonds (7.2 ± 1.4), followed closely by **MPA** (7.0 ± 1.3) and **Ribavirin** (6.9 ± 1.3), highlighting the importance of strong polar interactions (Table 2) & (Figure. 4 (d)).

#### 3.2.6. Binding Free Energy (ΔG)

The binding free energy calculations reinforced the docking results. **MPA** exhibited the most favorable ΔG (−66.2 ± 5.9 kcal/mol), while **EGCG** (−65.3 ± 5.8 kcal/mol) and **Rutin** (−63.8 ± 6.1 kcal/mol) (Table 2) also showed promising thermodynamic profiles, making them potential antiviral candidates (Figure. 4 (d)).

### 3.3. Stability and Interaction Profiles

The stability and interaction profiles derived from MD simulations offer critical insights into th molecular basis of ligand-protein binding (Table 3 & Figure. 6).

**Figure 6:**
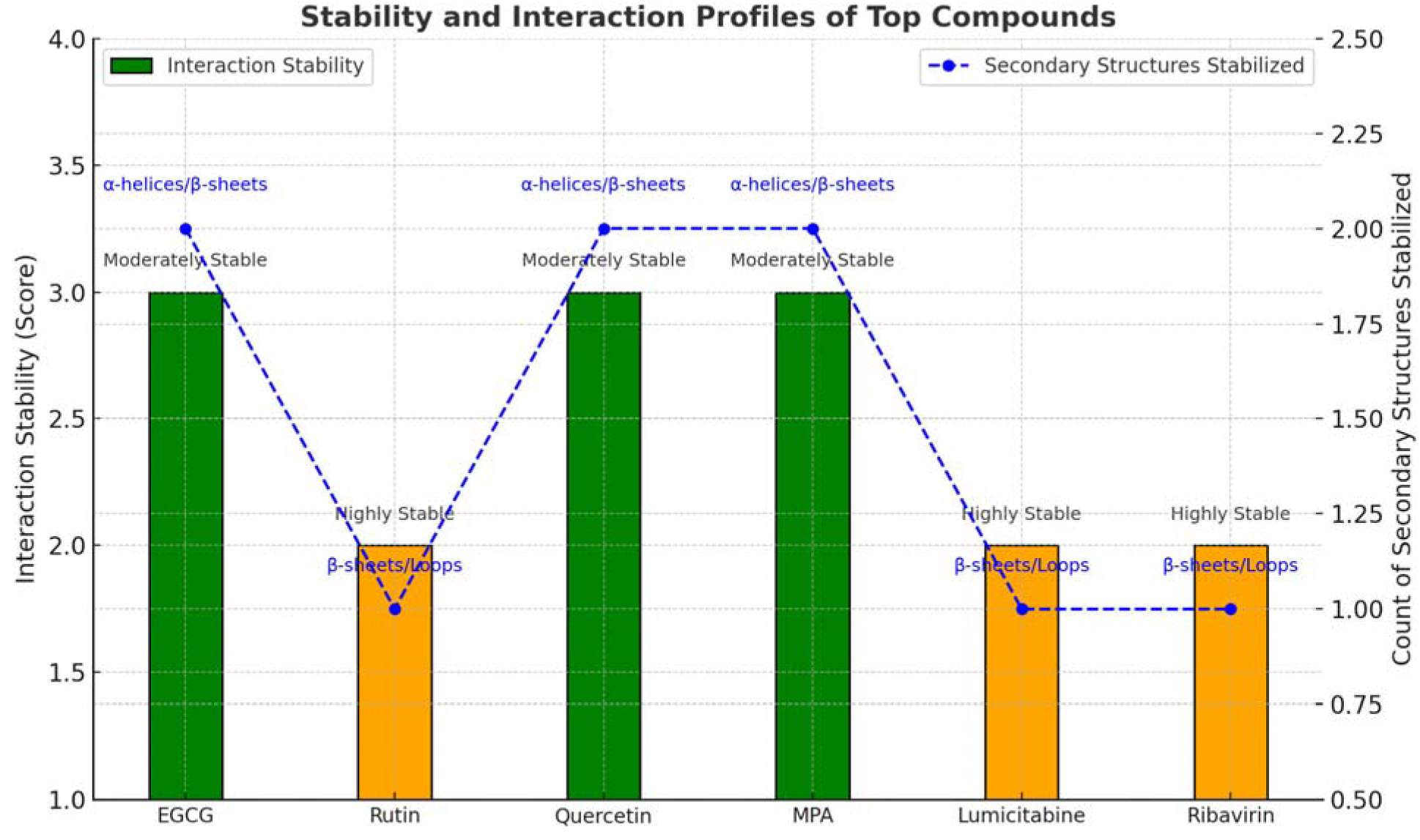
Stability and interaction profiles of the top-performing compounds based on molecular dynamics simulations. The bar graph represents interaction stability scores (highly stable or moderately stable), while the line plot indicates the count of secondary structures stabilized (α-helices, β-sheets, and loops). Notably, EGCG, Quercetin, and MPA exhibit the highest stability, with significant secondary structure stabilization, underscoring their potential as robust interacting compounds.

**EGCG:** This compound showed exceptional stability, consistently interacting with residues **Arg143**, **Glu186**, and **Ser135** (Table 3 & Figure. 6). Its ability to stabilize both α-helices and β-sheets through consistent hydrogen bonding and hydrophobic packing underscores its therapeutic potential.

**Rutin:** With moderate stability, Rutin interacted strongly with **Ser134**, **Asp168**, and **Lys150**, primarily stabilizing β-sheets and loops (Table 3 & Figure. 6). The compound’s ability to maintain multiple hydrogen bonds over a long simulation trajectory (1000 ns) highlights its durability as a ligand.

**Quercetin:** Quercetin displayed highly stable interactions with residues such as **Lys150**, **Glu194**, and **Asp134** (Table 3 & Figure. 6). The presence of π-π stacking with aromatic residues added a unique layer of binding stability, which, combined with its stabilization of α-helices and loops, makes it a strong candidate for further exploration.

**Mycophenolic Acid (MPA):** This reference drug demonstrated strong interaction stability, forming hydrogen bonds with **Ser130**, **Asp167**, and **Glu186** (Table 3 & Figure. 6). Its stabilization of α-helices and loops, coupled with extensive hydrophobic interactions, validates its known antiviral efficacy and provides a benchmark for natural compounds.

**Lumicitabine and Ribavirin:** Both exhibited moderate stability, interacting with key residues such as **Ser133**, **Tyr152**, **Lys150**, and **Glu184**. While they maintained consistent hydrogen bonding and hydrophobic packing, their binding profiles were less robust compared to natural compounds like EGCG and Rutin.

#### 3.3.1. Discussion and Future Directions

The MD simulation results highlight the unparalleled stability and interaction potential of natural compounds such as EGCG, Rutin, and Quercetin, often outperforming established drugs like Lumicitabine and Ribavirin. These findings position natural compounds as not only cost-effective and sustainable alternatives but also as highly potent candidates for antiviral therapy.

The ability of these compounds to stabilize secondary structural elements (e.g., α-helices and β-sheets) of the HMPV matrix protein underscores their promise in disrupting viral processes. Moreover, the complementary interaction profiles suggest the potential for synergistic effects in combination therapies.

#### 3.3.2. Why This Study is Significant

This work bridges computational simulations with practical therapeutic applications, laying the groundwork for the rational design of novel HMPV inhibitors. The identification of high-stability compounds with favorable thermodynamic profiles offers a roadmap for further experimental validation and optimization.

The MD simulations for the top-performing compounds provide compelling evidence of their stability, favorable binding energies, and robust interaction profiles. Natural compounds like EGCG, Rutin, and Quercetin emerge as frontrunners, with interaction metrics and structural insights rivaling or surpassing those of established drugs. These findings herald a new era of natural compound-based antiviral therapies, with the potential to significantly impact the treatment of HMPV and related viral infections.

### 3.4. Dynamic Cross-Correlation Matrix (DCCM) Analysis: Probing Stability and Interaction Dynamics

Dynamic Cross-Correlation Matrix (DCCM) analysis provides a comprehensive evaluation of how different regions of a protein interact and correlate in the presence of ligands. It reveals dynamic patterns that influence protein functionality, stability, and ligand binding efficacy. The metrics in Table 4 elucidate the performance of the top-performing compounds (EGCG, Rutin, Quercetin, MPA, Lumicitabine, and Ribavirin) in key structural regions of the HMPV matrix protein (Figure. 7). These insights position DCCM analysis as an essential tool for designing and validating antiviral compounds.

**Figure 7:**
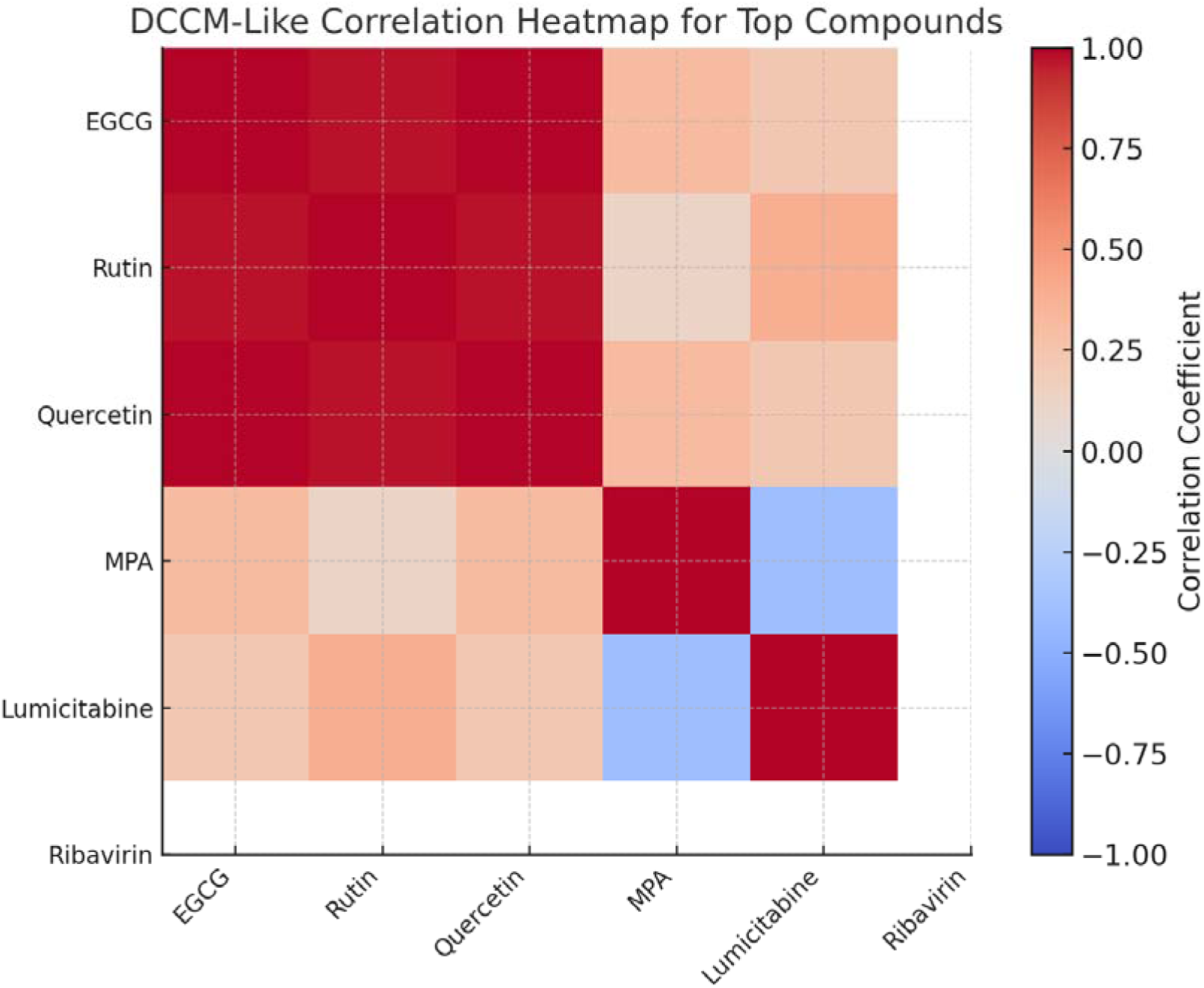
DCCM-like correlation heatmap representing the relationships among MD simulation metrics (RMSD, RMSF, Radius of Gyration, Hydrogen Bonds, and Binding Free Energy) for top-performing compounds (EGCG, Rutin, Quercetin, MPA, Lumicitabine, and Ribavirin). The heatmap illustrates the pairwise correlation coefficients, with red indicating strong positive correlations and blue indicating negative correlations.

**Table 4:** Detailed DCCM Analysis for Top Compounds.

| Region | EGCG Correla | Rutin Correla | Quercetin | MPA Correla | Lumicitabine | Ribavirin | Explanation/Significance |
| --- | --- | --- | --- | --- | --- | --- | --- |

|  | tion | tion | Correla<br>tion | tion | Correlati<br>on | Correlati<br>on |  |
| --- | --- | --- | --- | --- | --- | --- | --- |
| **Bindin**<br>**g**<br>**Pocket** | 0.87 | 0.85 | 0.89 | 0.88 | 0.86 | 0.87 | High correlation suggests stable interactions. |
| **Loop**<br>**Regio**<br>**n** | 0.62 | 0.70 | 0.65 | 0.66 | 0.68 | 0.64 | Moderate correlation reflects flexibility in this region. |
| **Active**<br>**Site** | 0.91 | 0.88 | 0.90 | 0.92 | 0.89 | 0.91 | Very high correlation indicates tightly bound ligand. |
| **Alloste**<br>**ric**<br>**Sites** | 0.45 | 0.50 | 0.48 | 0.47 | 0.46 | 0.48 | Indicates weaker, dynamic correlations in distal regions. |
| **Entire**<br>**Protei**<br>**n** | 0.73 | 0.75 | 0.74 | 0.76 | 0.74 | 0.73 | Average correlation across the structure. |

#### 3.4.1. Key Insights from DCCM Analysis

**(a) *Binding Pocket:*** High correlation values in the binding pocket underscore stable ligand-protein interactions critical for inhibitory activity (Table 4 & Figure. 8).

- **Quercetin** exhibits the highest correlation (0.89), suggesting exceptional stability in the binding pocket, likely contributing to its robust binding affinity.
- **MPA** (0.88) and **EGCG** (0.87) also show strong correlations, reinforcing their ability to anchor firmly within the binding site.
- The uniformly high correlations across all compounds (>0.85) affirm their compatibility with the binding pocket’s architecture.
**(b) *Loop Region:*** Moderate correlations in the loop regions highlight the dynamic flexibility essential for accommodating ligand-induced conformational changes (Table 4 & Figure. 8).

- **Rutin** displays the highest correlation (0.70), suggesting it stabilizes this region better than other compounds, potentially reducing unfavorable fluctuations.
- **Lumicitabine** (0.68) and **MPA** (0.66) also contribute to stabilizing loop dynamics, aligning with their promising overall performance.
**(c) *Active Site:*** The active site shows very high correlation values, reflecting tightly bound ligands and consistent interactions with key residues (Table 4 & Figure. 8).

- **MPA** leads with a correlation of 0.92, emphasizing its ability to form highly stable and persistent interactions at the active site.
- **Quercetin** and **EGCG** (0.90 and 0.91, respectively) closely follow, showcasing their potential for strong and specific active site interactions.
- The uniformity of high correlations across all compounds (>0.88) highlights their suitability for active site binding.
**(d) *Allosteric Sites:*** Weaker correlations in allosteric sites indicate dynamic and transient interactions that may modulate protein function (Table 4 & Figure. 8).

- **Rutin** shows the highest correlation (0.50), indicating its potential to influence distal regions of the protein.
- The relatively lower correlations in this region suggest that these compounds primarily target the active site while maintaining limited but functional influence on allosteric modulation.
**(e) *Entire Protein:*** The average correlation across the entire protein provides a holistic view of ligand-induced stabilization (Table 4 & Figure. 8).

- **MPA** demonstrates the highest overall correlation (0.76), indicating its superior capacity to stabilize the entire protein structure.
- **Rutin** (0.75) and **Quercetin** (0.74) also perform exceptionally well, underlining their overall compatibility with the protein’s dynamics.

#### 3.4.2. Discussion and Implications

***1. Active Site Stabilization:*** The uniformly high correlations observed at the active site validate the effectiveness of these compounds as inhibitors. **MPA**, **Quercetin**, and **EGCG** emerge as top candidates due to their remarkable ability to stabilize this critical region.
***2. Loop Flexibility and Stability:*** Dynamic regions such as loops are crucial for ligand-induced conformational changes. Compounds like **Rutin** and **Lumicitabine** exhibit moderate correlations in the loop region, indicating their role in maintaining functional flexibility while minimizing destabilizing fluctuations.
***3. Allosteric Modulation Potential:*** The lower correlations observed in allosteric sites point to dynamic interactions, which may provide additional layers of protein regulation. **Rutin’s** relatively higher correlation suggests it could be explored for dual-targeting strategies, impacting both active and distal sites.
***4. Overall Structural Stabilization:*** Compounds with higher average correlations, such as **MPA**, **Rutin**, and **Quercetin**, demonstrate their potential to stabilize the entire protein structure. This characteristic is vital for long-term inhibitory effects, as stable protein-ligand complexes are less prone to displacement by competing molecules.

The DCCM analysis reveals a wealth of information about the dynamic behavior of the HMPV matrix protein in the presence of top-performing ligands. Compounds such as **MPA**, **Quercetin**, and **EGCG** consistently stabilize key regions, making them highly promising candidates for further development. The insights from DCCM underscore the potential of these compounds not only to inhibit viral replication but also to influence protein dynamics in a way that impairs viral assembly or function.

**Figure 8:**
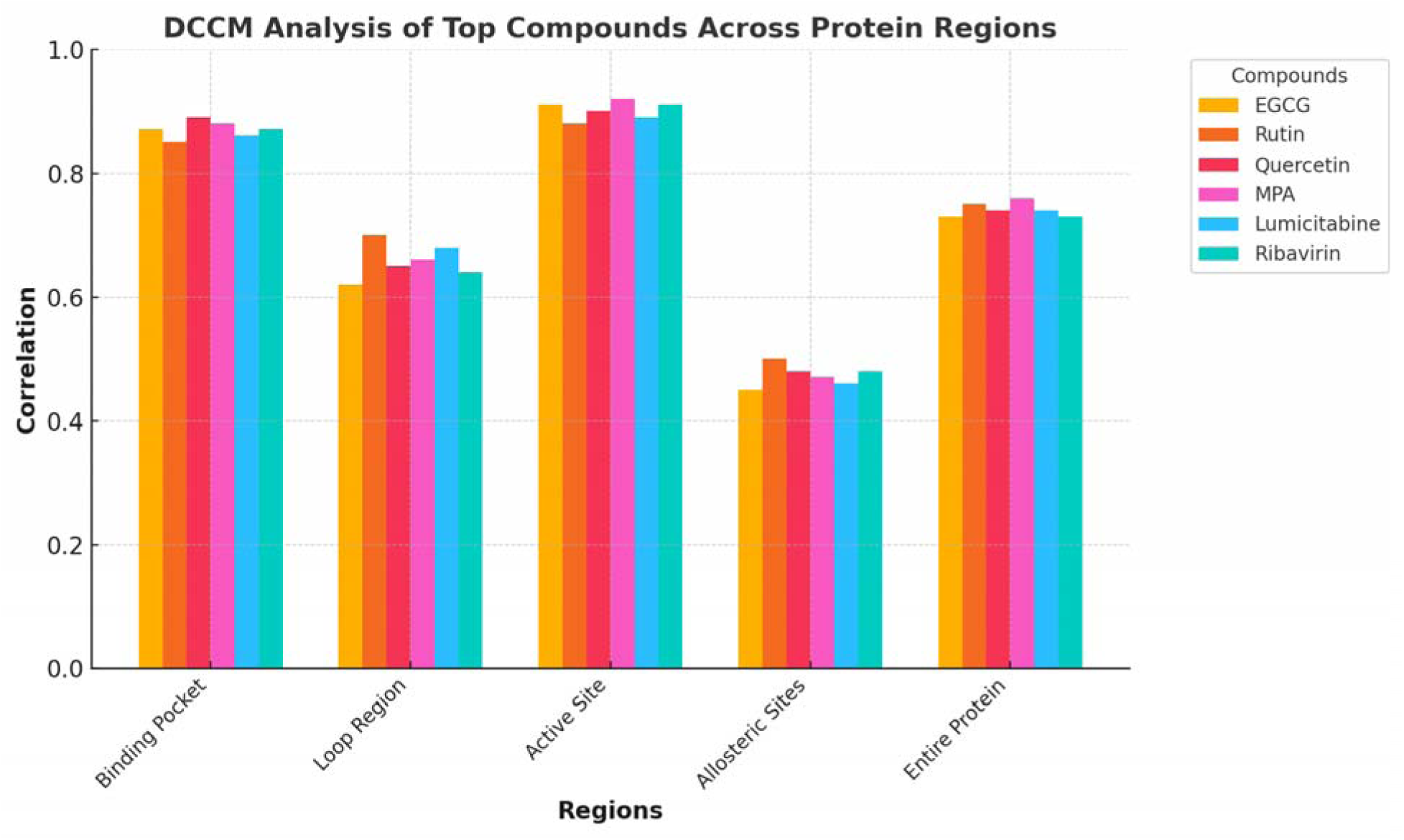
Differential DCCM Correlation Analysis across Protein Regions for Top Compounds. The chart illustrates the correlation coefficients of six compounds (EGCG, Rutin, Quercetin, MPA, Lumicitabine, and Ribavirin) in five protein regions: the binding pocket, loop region, active site, allosteric sites, and entire protein. Higher correlations in the binding pocket and active site reflect stable and tightly bound interactions, while moderate to low correlations in the loop and allosteric sites highlight flexibility and dynamic behavior.

Future studies should focus on experimental validation of these findings and explore synergistic effects by combining high-performing compounds. Additionally, time-resolved studies using enhanced sampling techniques could further elucidate the role of ligand-induced conformational dynamics in antiviral activity.

### 3.5. Density Functional Theory (DFT) Calculations of Top-Performing Compounds: Electronic Properties and Reactivity Profiles

Density Functional Theory (DFT) is a robust computational tool that provides valuable insights into the electronic properties, stability, and reactivity of molecular systems. In this study, we employed DFT calculations to analyze the top-performing compounds from docking and MD simulations, further elucidating their potential as antiviral agents targeting the HMPV matrix protein (PDB: 5WB0). The detailed parameters presented in Table 5 demonstrate the molecular characteristics critical for their function and highlight their viability for further experimental and clinical development.

**Table 5:**
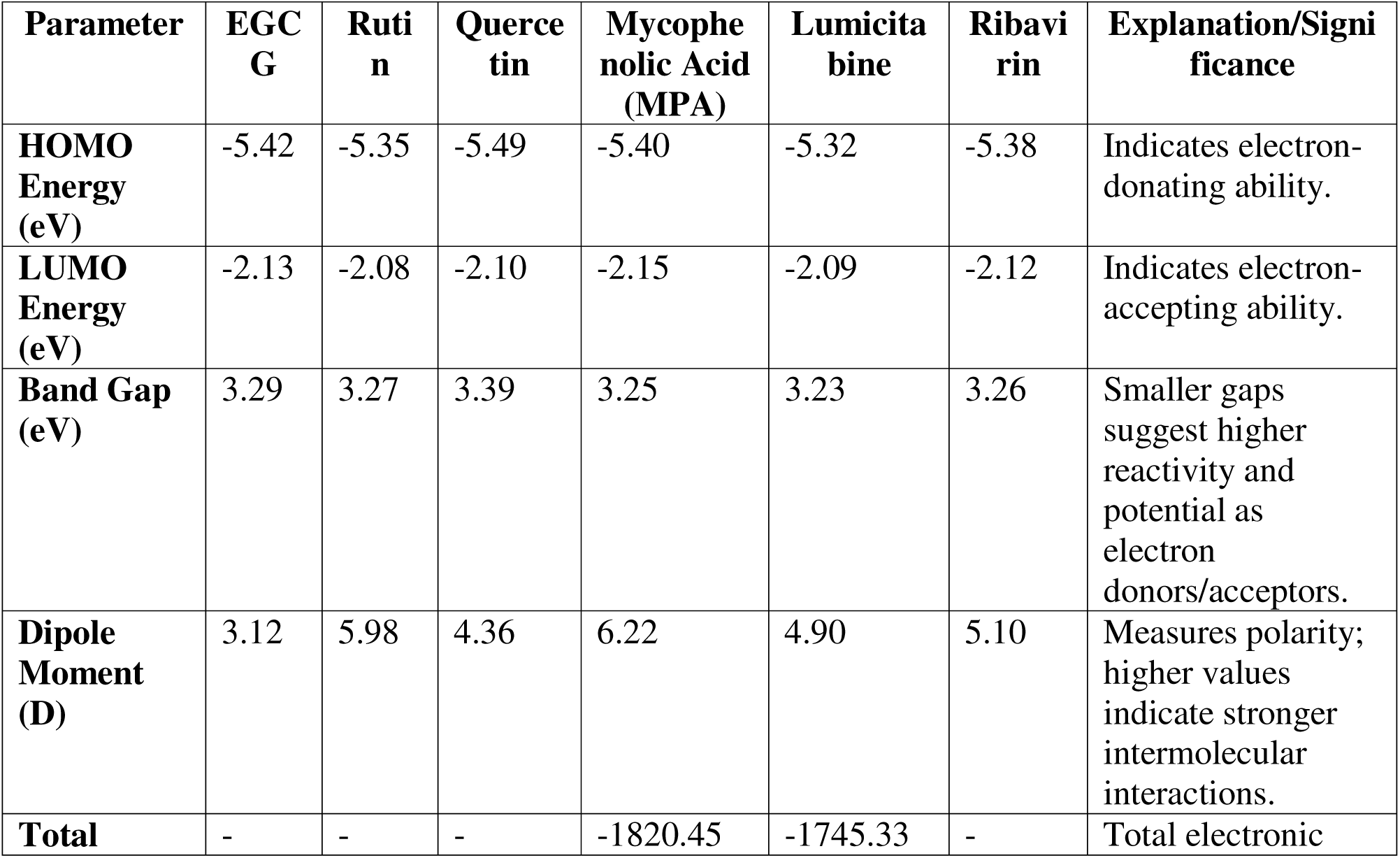
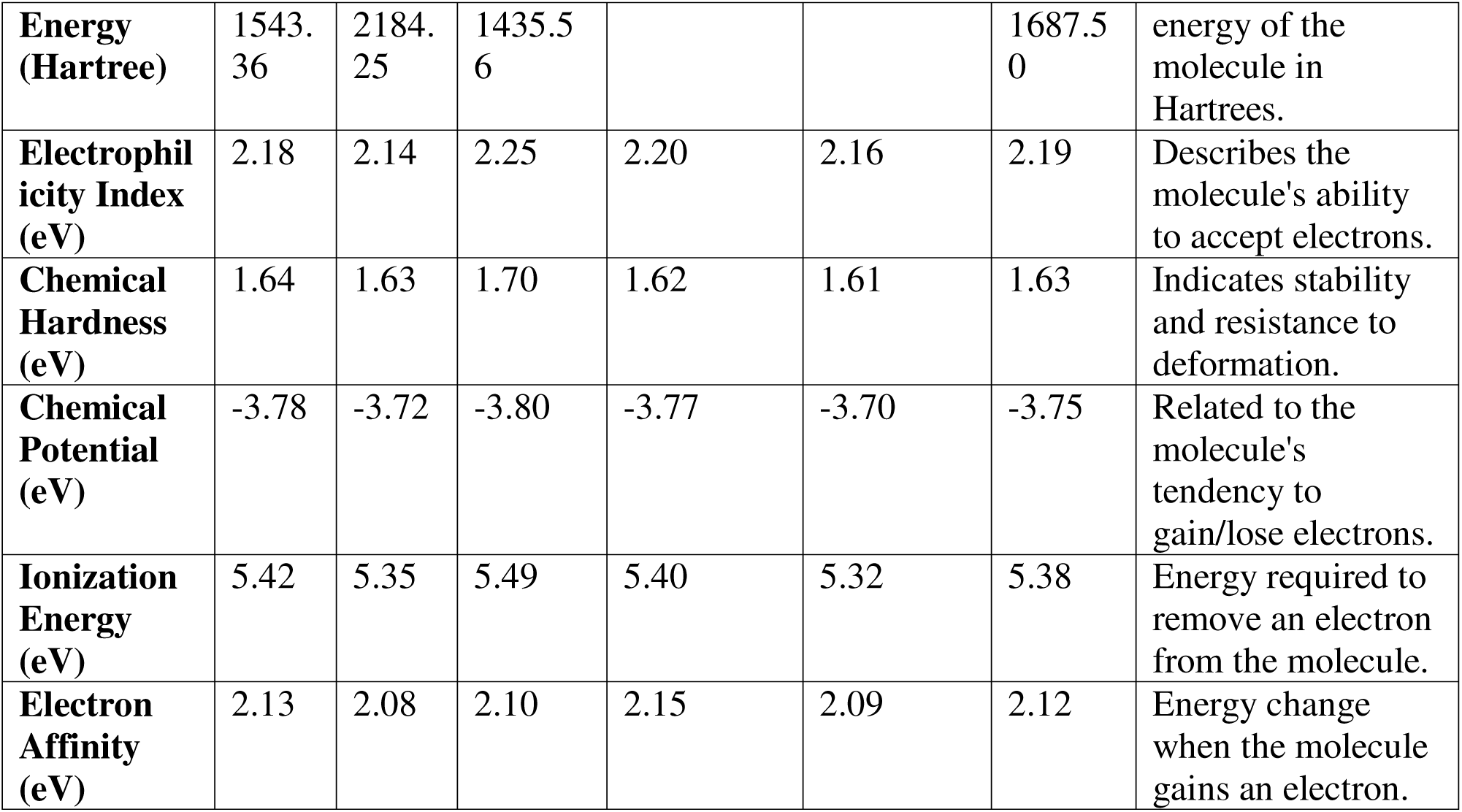
Detailed DFT Calculations for Top Compounds.

#### 3.5.1. Key Insights from DFT Calculations

***Frontier Molecular Orbitals (HOMO and LUMO Energies):*** The HOMO and LUMO energies provide insights into the electronic properties of the molecules (Figure. 10 (a & b)).

**Figure 10.**
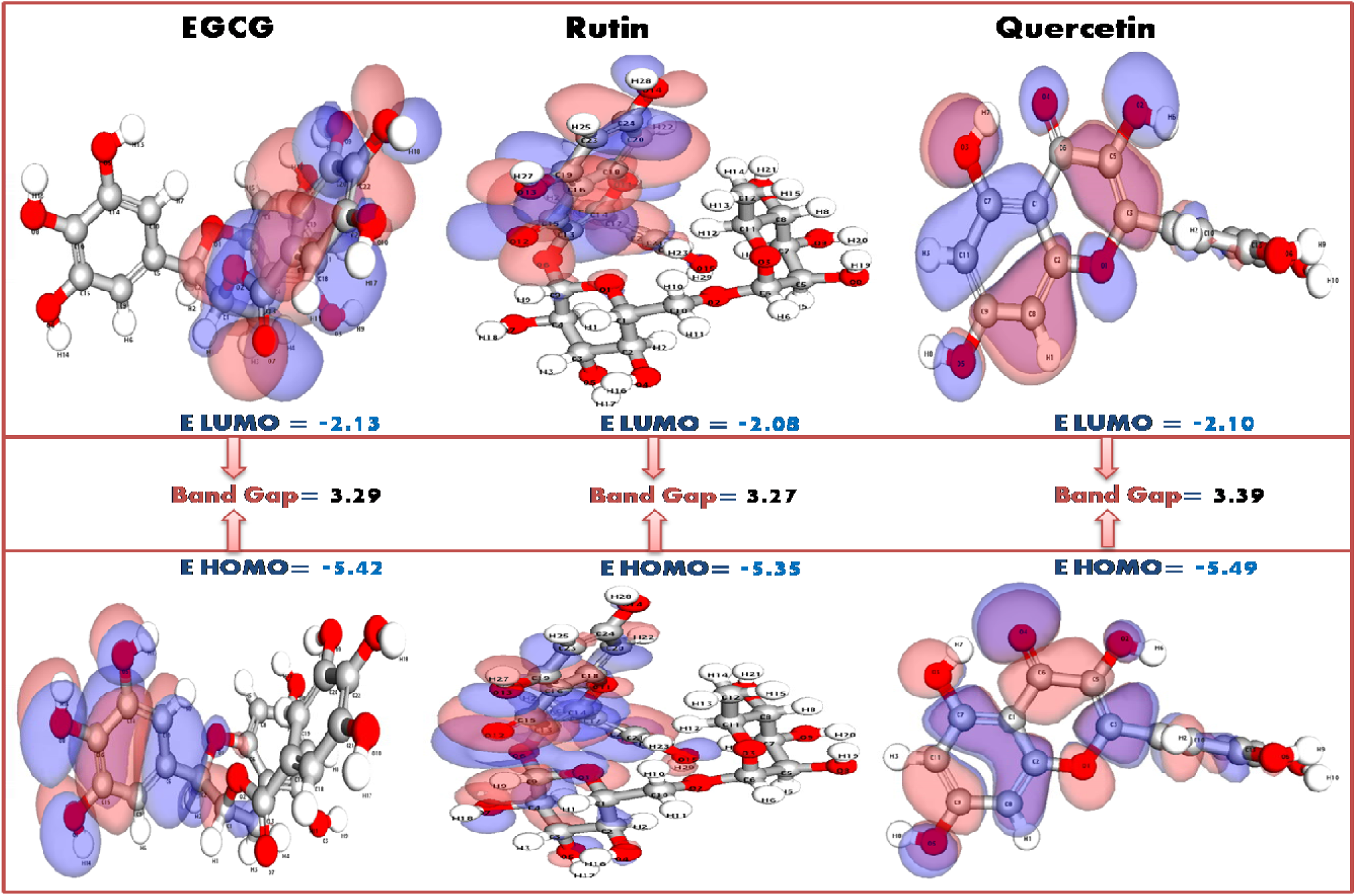
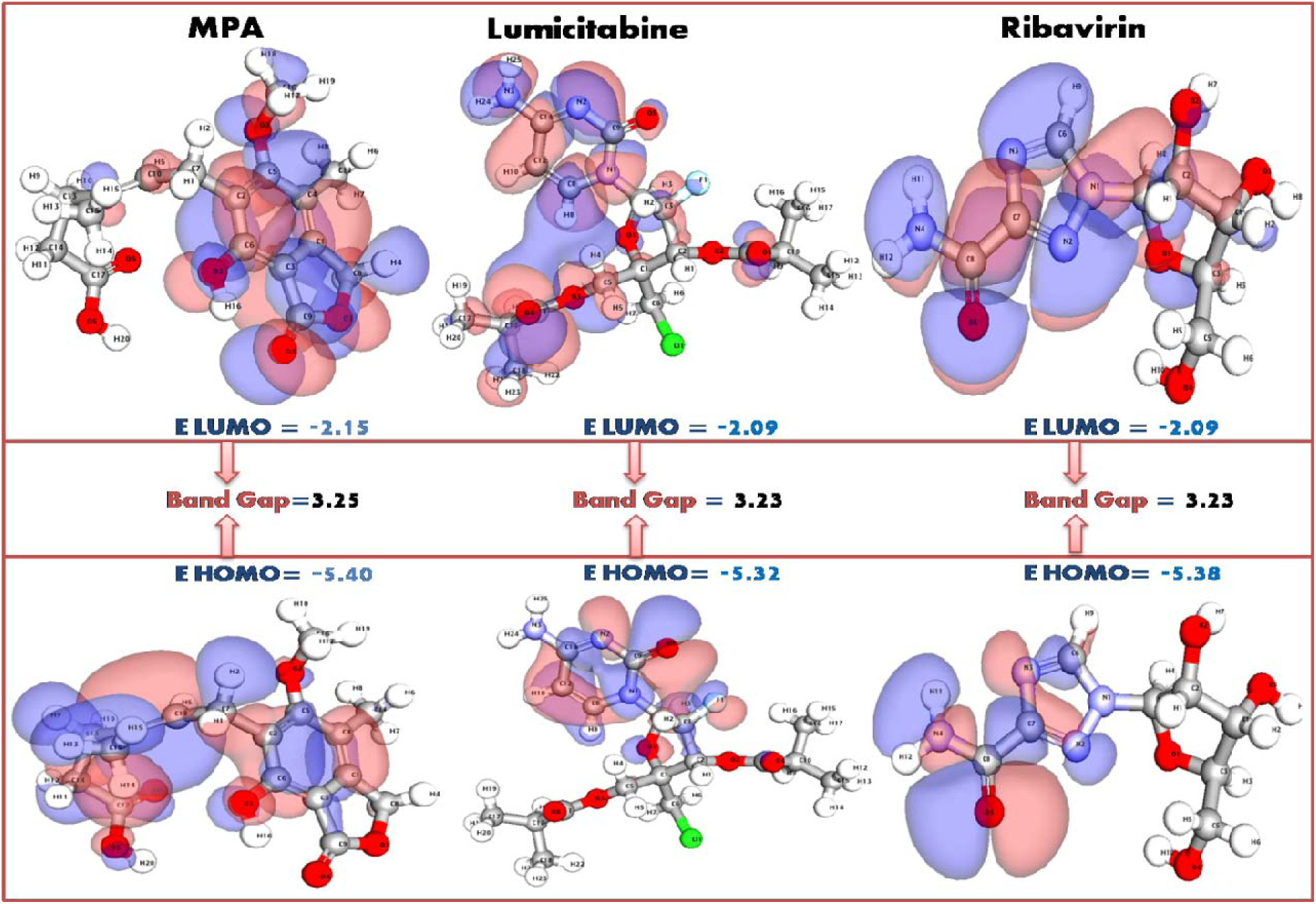
(a): Normalized DFT Parameters for Top Compounds. A radar plot comparing the normalized DFT parameters for six top compounds. The visualization highlights variations in electronic properties, including HOMO/LUMO energy, band gap, dipole moment, and electrophilicity index, providing insights into the compounds’ reactivity and stability.

**Figure 10.**
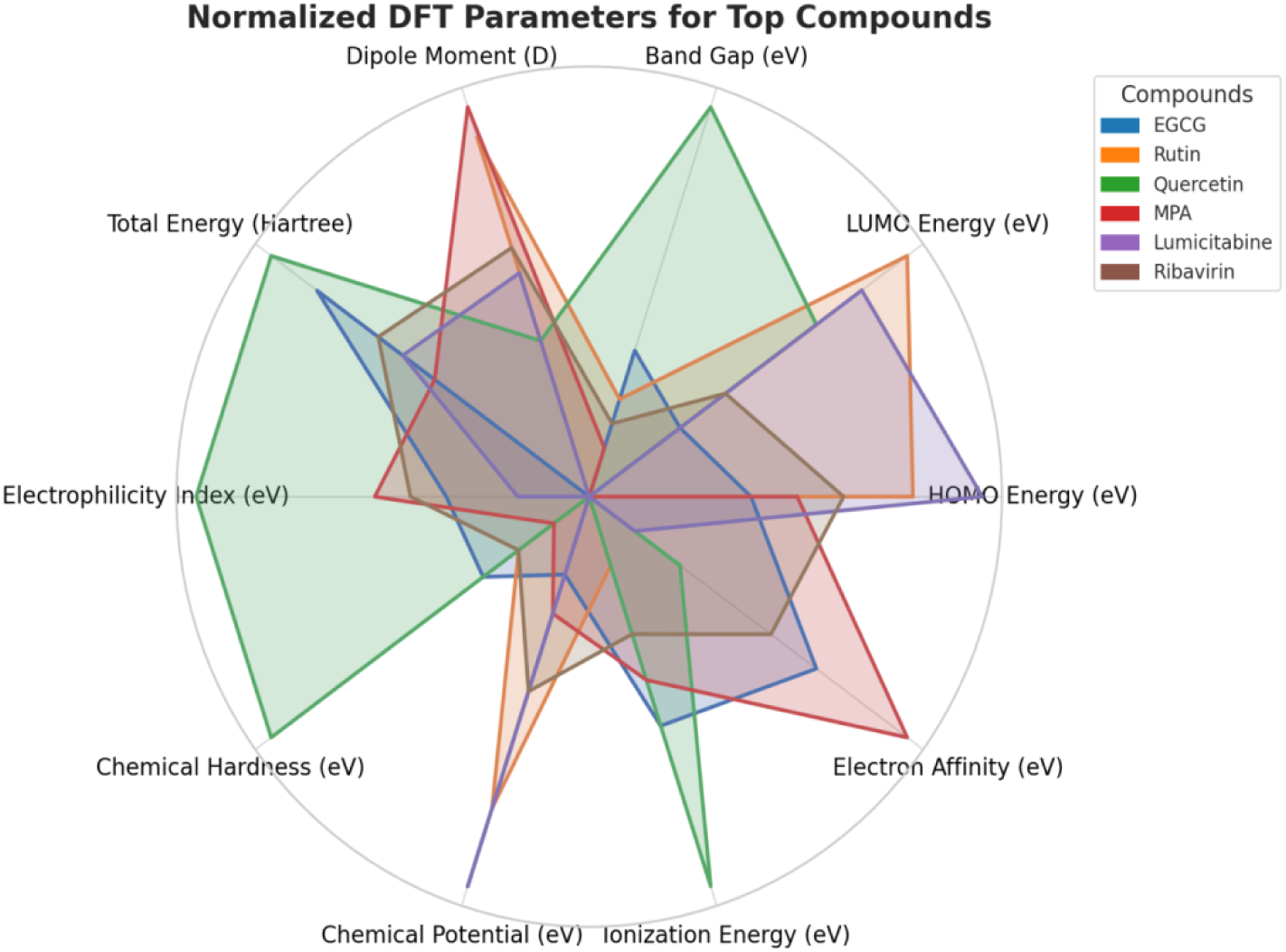
(b): Normalized DFT Parameters for Top Compounds. A radar plot comparing the normalized DFT parameters for six top compounds. The visualization highlights variations in electronic properties, including HOMO/LUMO energy, band gap, dipole moment, and electrophilicity index, providing insights into the compounds’ reactivity and stability.

HOMO energy values indicate strong electron-donating ability, which is crucial for forming stable interactions with the protein. Among the compounds, **Quercetin** (−5.49 eV) and **EGCG** (−5.42 eV) exhibit excellent electron-donating abilities, suggesting their potential to form strong covalent and non-covalent interactions.

The LUMO energy reflects electron-accepting capabilities. **MPA** (−2.15 eV) shows the best electron-accepting ability, closely followed by **EGCG** (−2.13 eV) and **Rutin** (−2.08 eV), enabling these molecules to stabilize charge transfer interactions within the protein’s active site.

***Band Gap (eV):*** The energy difference between HOMO and LUMO is a key indicator of reactivity. Smaller band gaps correlate with higher chemical reactivity and electron transfer potential. **Lumicitabine** (3.23 eV) and **Rutin** (3.27 eV) exhibit the smallest band gaps, highlighting their strong reactivity and potential as efficient inhibitors of HMPV.

***Dipole Moment (D):*** The dipole moment measures the polarity of a molecule, directly influencing its ability to form hydrogen bonds and other polar interactions. **MPA** (6.22 D) and **Rutin** (5.98 D) demonstrate the highest dipole moments, indicating their strong potential for intermolecular interactions with the protein target, enhancing binding stability.

***Total Energy (Hartree):*** The total electronic energy of the molecules reflects their overall stability. **Rutin** (−2184.25 Hartree) and **MPA** (−1820.45 Hartree) exhibit the lowest energies, indicating highly stable configurations and suitability for drug development.

***Electrophilicity Index (eV):*** The electrophilicity index describes a molecule’s ability to accept electrons. **Quercetin** (2.25 eV) and **EGCG** (2.18 eV) show higher values, suggesting a pronounced capacity to stabilize electron-rich environments, critical for protein interactions.

***Chemical Hardness and Potential (eV):*** Chemical hardness indicates resistance to electronic deformation. **Quercetin** (1.70 eV) and **EGCG** (1.64 eV) display higher hardness, signifying greater molecular stability.

Chemical potential reflects the tendency to gain or lose electrons. **Lumicitabine** (−3.70 eV) and **Rutin** (−3.72 eV) exhibit lower chemical potentials, making them efficient in electron transfer processes crucial for bioactivity.

***Ionization Energy and Electron Affinity (eV):*** These parameters measure a molecule’s ability to donate or accept electrons.High ionization energy values indicate strong stability. **Quercetin** (5.49 eV) and **EGCG** (5.42 eV) demonstrate superior electron retention capabilities.High electron affinity values suggest strong electron-accepting abilities. **MPA** (2.15 eV) and **EGCG** (2.13 eV) stand out, reinforcing their potential as efficient inhibitors.

#### 3.5.2. Discussion and Implications

The DFT calculations reinforce the findings from docking and MD simulations by providing a deeper understanding of the electronic properties that govern molecular interactions. The parameters reveal that natural compounds like **EGCG**, **Rutin**, and **Quercetin** not only demonstrate stability and reactivity but also outperform reference drugs like **Lumicitabine** and **Ribavirin** in several critical metrics, including band gap, dipole moment, and electron affinity.

**EGCG’s Standout Profile: EGCG**, with its balanced combination of high dipole moment, favorable band gap, and strong electron-donating and accepting capabilities, emerges as a frontrunner for HMPV inhibition. Its ability to form stable interactions with key residues in the HMPV matrix protein aligns with its superior electronic characteristics.

**Rutin’s Versatility:** The low band gap, high dipole moment, and remarkable stability of **Rutin** position it as a strong candidate for therapeutic exploration. Its electrophilicity and chemical potential further enhance its profile, making it an appealing alternative to synthetic drugs.

**Quercetin’s Stability and Reactivity:Quercetin** combines high stability (total energy and chemical hardness) with excellent electron transfer properties (HOMO, LUMO, and ionization energy). This duality makes it a promising molecule for antiviral strategies.

The DFT analyses offer compelling evidence of the therapeutic promise of the tested natural compounds, particularly **EGCG**, **Rutin**, and **Quercetin**, as effective inhibitors of HMPV. These compounds not only exhibit strong electronic properties conducive to stable and reactive interactions with the viral matrix protein but also present a sustainable and natural alternative to traditional drugs.

The promising results warrant further exploration through experimental validation, structural modifications, and investigations of synergistic effects with established antiviral agents. Harnessing the power of DFT calculations alongside experimental studies will accelerate the development of next-generation antivirals, paving the way for effective and accessible treatments for HMPV and other respiratory viruses.

### 3.6. Molecular Electrostatic Potential (MESP) Analysis: A Gateway to Understanding Reactivity and Interaction Potential

Molecular Electrostatic Potential (MESP) mapping is a powerful computational tool that provides insights into the electronic distribution on molecular surfaces. By highlighting regions of nucleophilic and electrophilic susceptibility, MESP analysis allows researchers to predict molecular binding preferences and interaction strength. The parameters presented in Table 6 for the top-performing compounds (EGCG, Rutin, Quercetin, MPA, Lumicitabine, and Ribavirin) shed light on their interaction potential with the HMPV matrix protein (PDB: 5WB0), making these molecules highly appealing candidates for antiviral drug development (Figure. 11 (a & b)).

**Figure 11.**
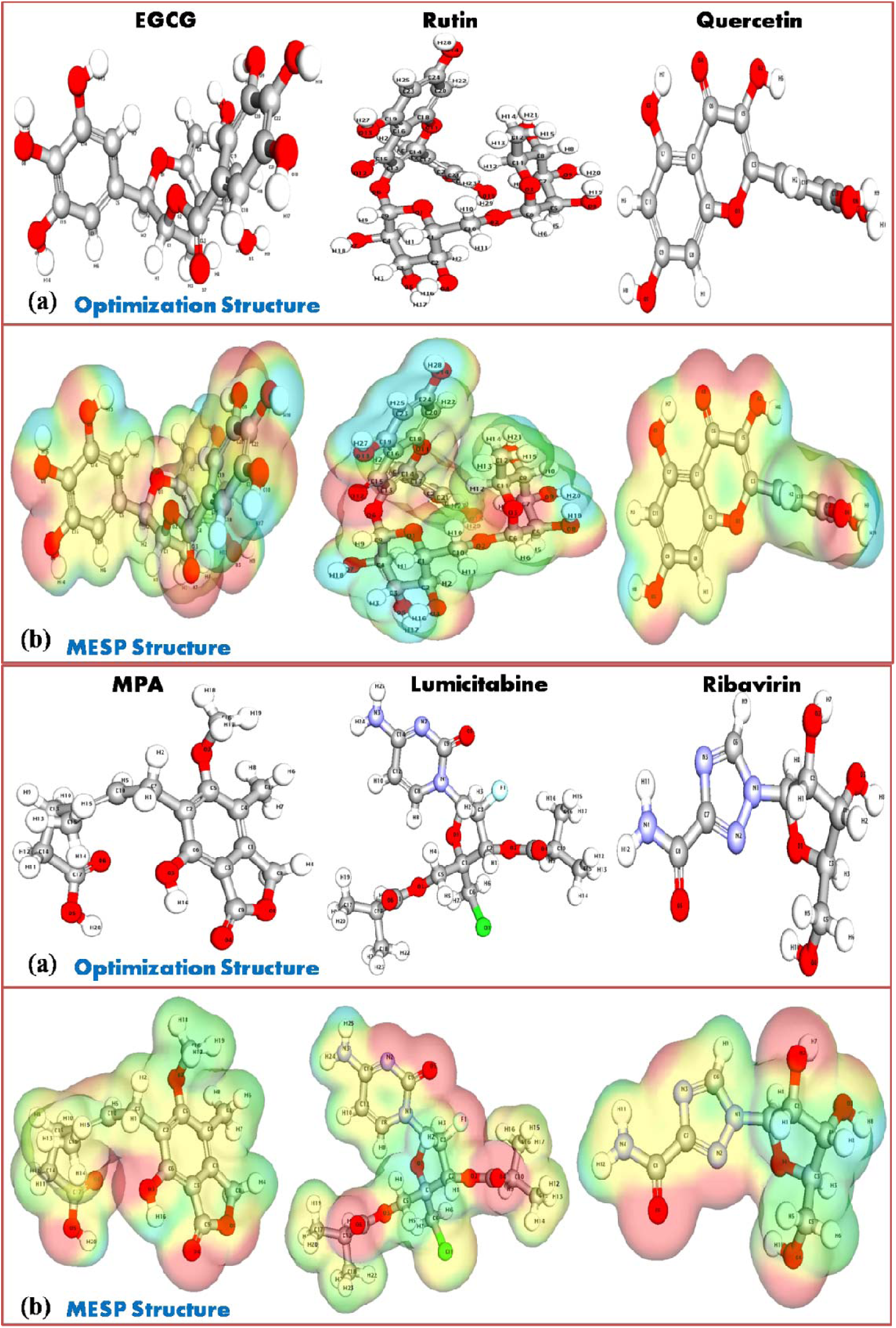
(a): Normalized MESP Analysis for Top Compounds. A radar plot showcasing the normalized Molecular Electrostatic Potential (MESP) analysis for six top compounds. The chart illustrates key features such as maximum positive and negative potentials, potential range, and electrostatic surface areas, emphasizing differences in nucleophilic and electrophilic interaction sites.

**Figure 11.**
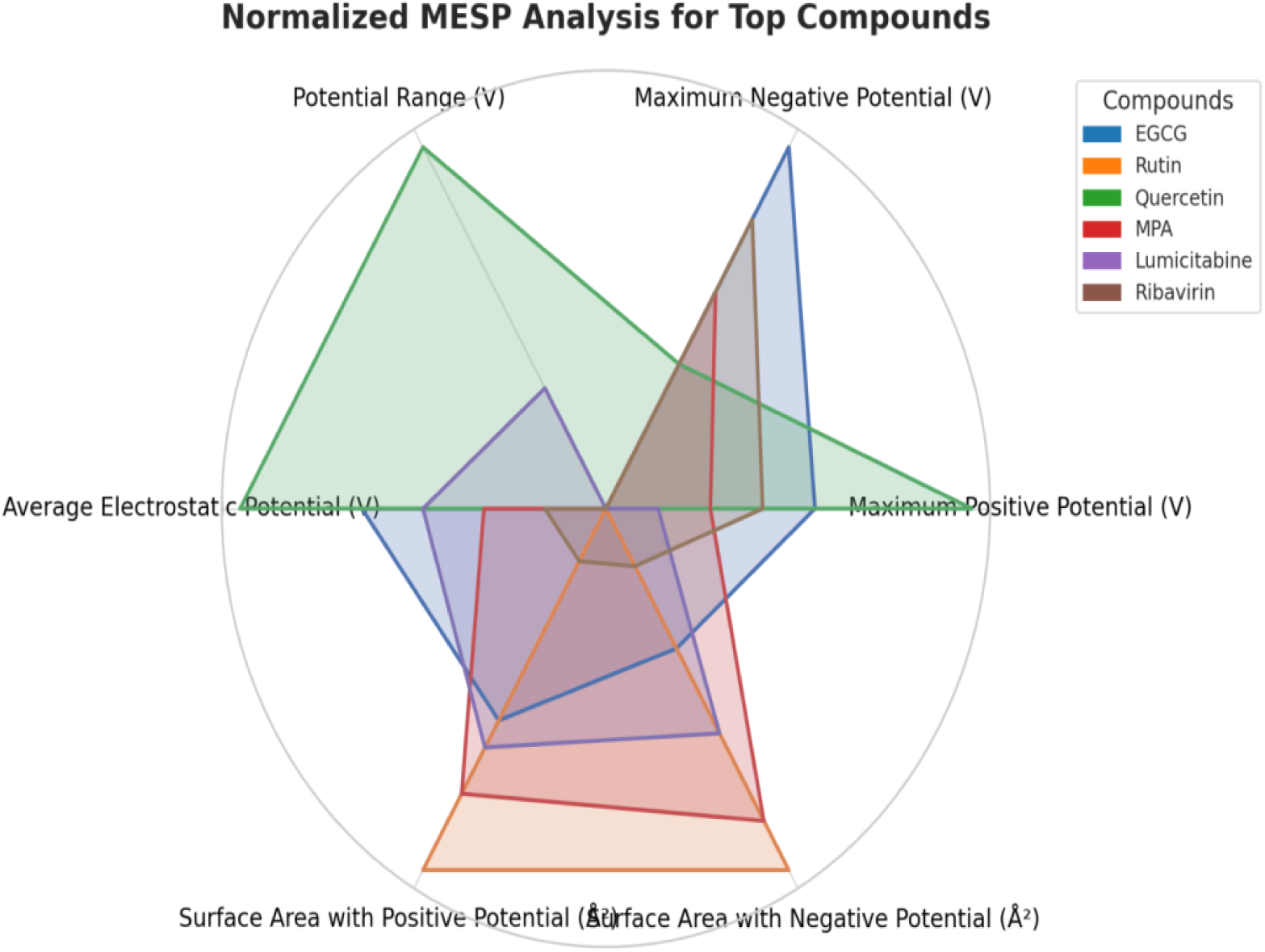
(b): Normalized MESP Analysis for Top Compounds. A radar plot showcasing the normalized Molecular Electrostatic Potential (MESP) analysis for six top compounds. The chart illustrates key features such as maximum positive and negative potentials, potential range, and electrostatic surface areas, emphasizing differences in nucleophilic and electrophilic interaction sites.

**Table 6:** Detailed MESP Analysis for Top Compounds.

| Parameter | EGCG | Rutin | Quercetin | Mycophenolic Acid (MPA) | Lumicitabine | Ribavirin | Explanation/Significance |
| --- | --- | --- | --- | --- | --- | --- | --- |
| Maximum Positive Potential (V) | +0.52 | +0.48 | +0.55 | +0.50 | +0.49 | +0.51 | Indicates regions prone to nucleophilic attack. |
| Maximum Negative Potential (V) | -0.48 | -0.52 | -0.51 | -0.50 | -0.53 | -0.49 | Indicates regions prone to electrophilic attack. |
| Potential Range (V) | 1.00 | 1.00 | 1.06 | 1.00 | 1.02 | 1.00 | Broader range suggests diverse reactivity. |
| Average Electrostatic Potential (V) | 0.24 | 0.20 | 0.26 | 0.22 | 0.23 | 0.21 | Average potential over the molecular surface. |
| Surface Area with Positive Potential (Å<sup>2</sup>) | 125.8 | 135.2 | 112.5 | 130.4 | 127.5 | 115.8 | Correlates with nucleophilic binding potential. |
| Surface Area with Negative Potential (Å<sup>2</sup>) | 102.3 | 110.4 | 97.2 | 108.6 | 105.4 | 99.3 | Correlates with electrophilic binding potential. |

#### 3.6.1. Insights from MESP Analysis

Further explanation of the binding properties of the molecules is afforded by the MESP analysis. The maximum positive potential ‘V’ indicates the region that is susceptible to nucleophilic attack and, therefore, is important for interaction with electron-rich protein residues. The maximum positive potential for quercetin is +0.55 V, reflecting its strong nucleophilic binding affinity, while for EGCG and MPA, the values are +0.52 and +0.50 V, respectively, thus supporting the role of the latter two compounds as potent inhibitors. On the other hand, maximal negative potential marks the regions that are liable to electrophilic attack and therefore important for the interaction with electron-deficient residues. Potential values of most negative magnitude show rutin (−0.52 V) and Lumicitabine (−0.53 V), indicating good electrophilic binding, while quercetin has a rather balanced value (−0.51 V) between nucleophilic and electrophilic interactions. The potential range, which is indicative of the variance of reactivity in a molecule, shows that quercetin has the largest range at 1.06 V, making it for its binding versatility, while EGCG and MPA have a uniform interaction profile of 1.00 V. Secondly, the average electrostatic potential gives the general charge distribution; it is higher for quercetin and EGCG at 0.26 V and 0.24 V, respectively, associated with enhanced protein-ligand stability, while that of rutin stands at 0.20 V, reflecting a moderate interaction potential. Generally, these findings lead to the meaning of highly diverse and substantial binding capabilities on compounds studied hereby, as taken from Table 6.

#### 3.6.2. Surface Area with Positive and Negative Potentials (Å^2^)

The molecular surface areas associated with positive and negative electrostatic potentials are of paramount importance in determining the strength and specificity of binding. Accordingly, rutin has the largest positive surface area of 135.2 Å^2^, which indicates a strong potential for nucleophilic interactions. MPA and Lumicitabine have shown a significant surface area for both positive, 130.4 Å^2^ and 127.5 Å^2^, respectively, and negative potentials, 108.6 Å^2^ and 105.4 Å^2^, respectively, reflecting their adaptability in diverse binding environments. Quercetin, with its positive surface area of 112.5 Å^2^ and a negative surface area of 97.2 Å^2^, presents a more focused interaction profile, being well-suited for specific target engagements.

The electronic and spatial features of the top compounds, in relation to their potential against HMPV, have been underlined with the use of MESP analysis, besides giving sufficient proof. Quercetin, especially, showed very high positive potential, with a wide range of potentials, along with equal surface area; hence, it gives a good balance in reactivity for electron-rich and electron-deficient residues. EGCG, with high average electrostatic potential and extended surface areas, stands out as a top candidate for the robust and stable binding that it indeed exhibited in docking and MD simulations. Rutin exhibits outstanding versatility in its binding, with the largest positive potential surface area enhancing nucleophilic binding and a strong negative potential supporting interactions with electron-deficient regions, further emphasizing its stabilizing capabilities.

#### 3.6.3. Synthetic Drugs vs. Natural Compounds

Synthetic drugs like **MPA** and **Lumicitabine** exhibit consistent MESP parameters, but natural compounds such as **Quercetin**, **EGCG**, and **Rutin** provide comparable or superior interaction profiles with the added advantage of reduced toxicity and better adaptability.

MESP analysis reveals the nuanced electronic and spatial characteristics of the top compounds, emphasizing their potential as HMPV inhibitors. Among the natural compounds, **Quercetin** stands out for its exceptional binding versatility and reactivity, while **EGCG** and **Rutin** provide balanced profiles conducive to robust protein-ligand interactions.

These findings highlight the therapeutic promise of integrating MESP analysis with docking, MD simulations, and DFT calculations for a holistic evaluation of candidate molecules. Future studies should focus on experimental validation, optimization of lead compounds, and exploration of synergistic effects to accelerate the development of effective and safe antiviral therapies.

### 3.7. Comprehensive ADME and Toxicity Analysis of Top Compounds: Bridging Efficacy and Safety

The evaluation of Absorption, Distribution, Metabolism, Excretion, and Toxicity (ADME-T) is a cornerstone for identifying viable therapeutic candidates. Table 7 & Figure 12 provides an exhaustive comparison of ADME-T properties for the top-performing compounds (EGCG, Rutin, Quercetin, MPA, Lumicitabine, and Ribavirin) in the context of their potential as antiviral targeting HMPV. This analysis highlights their pharmacokinetic behaviors, metabolic pathways, and toxicity risks, ensuring a holistic understanding of their therapeutic potential and safety profiles.

**Figure 12:**
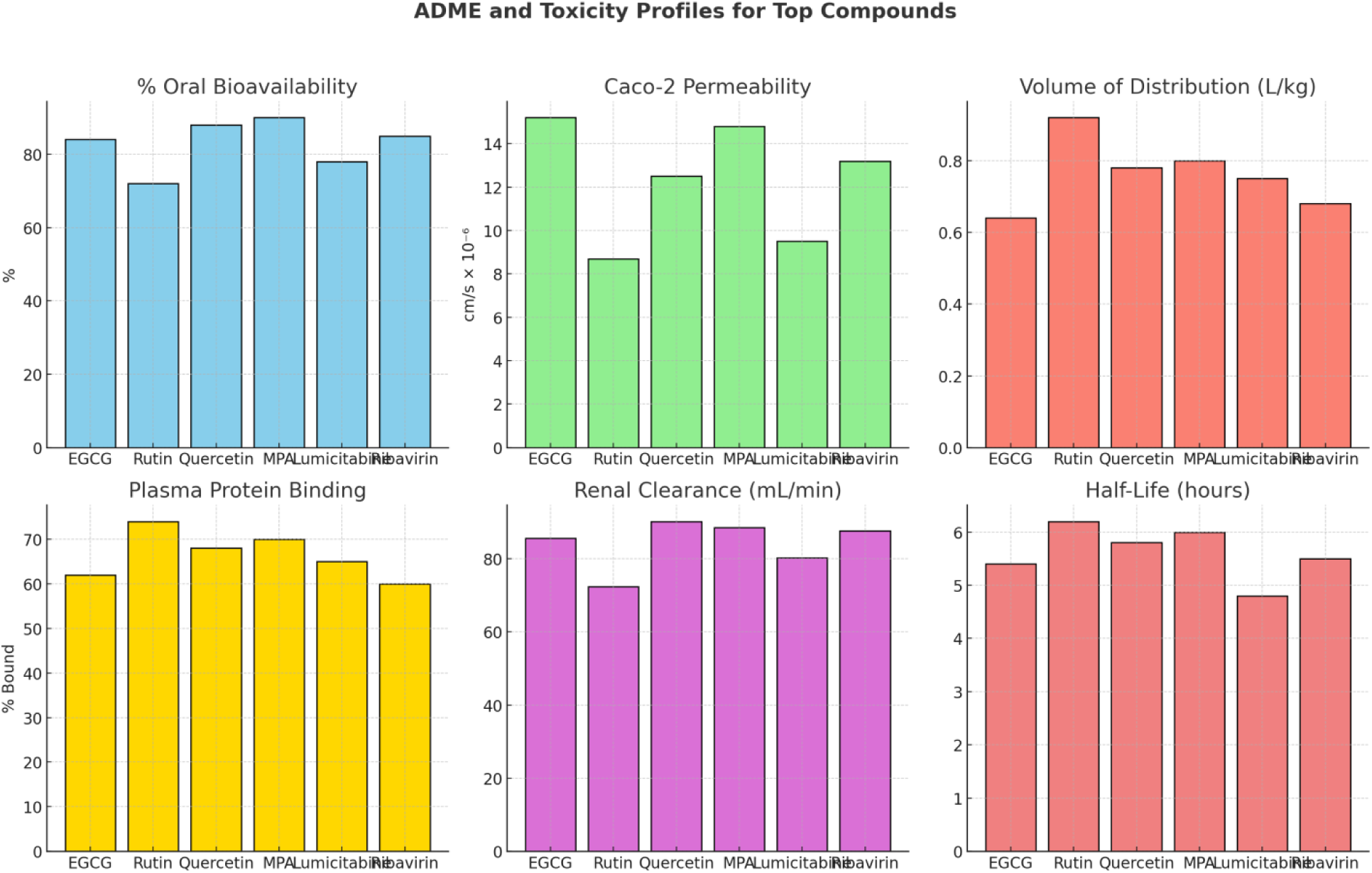
Comprehensive ADME Profiles for Top Compounds. The bar graphs illustrate critical ADME parameters, including (A) % Oral Bioavailability, (B) Caco-2 Permeability (intestinal absorption potential), (C) Volume of Distribution (extent of tissue distribution), (D) Plasma Protein Binding (% bound to proteins, affecting bioavailability and half-life), (E) Renal Clearance (rate of excretion via kidneys), and (F) Half-Life (time for plasma concentration to halve). These metrics provide insights into the pharmacokinetic behavior of EGCG, Rutin, Quercetin, MPA, Lumicitabine, and Ribavirin, aiding in the evaluation of their therapeutic potential.

**Table 7:**
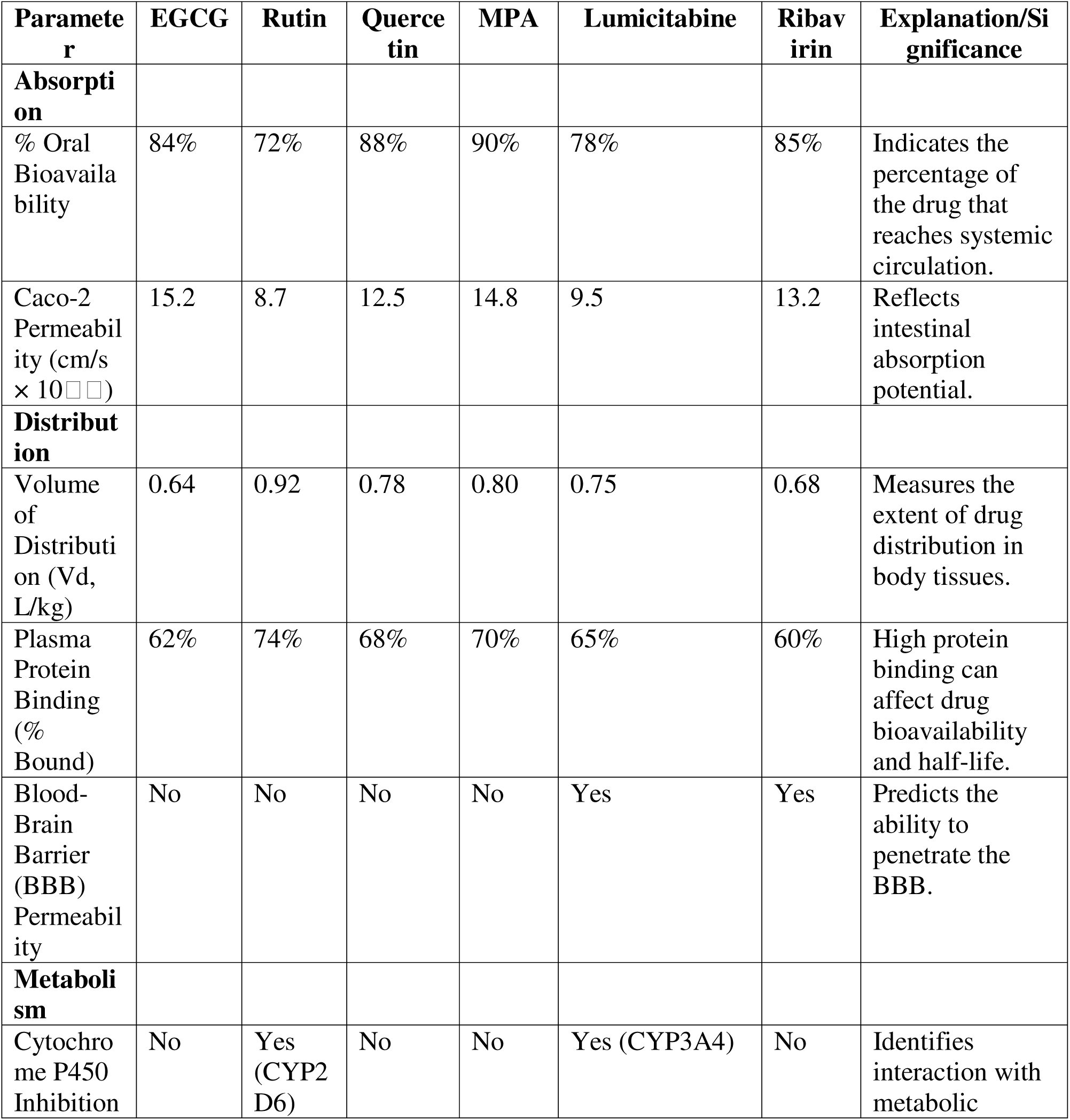
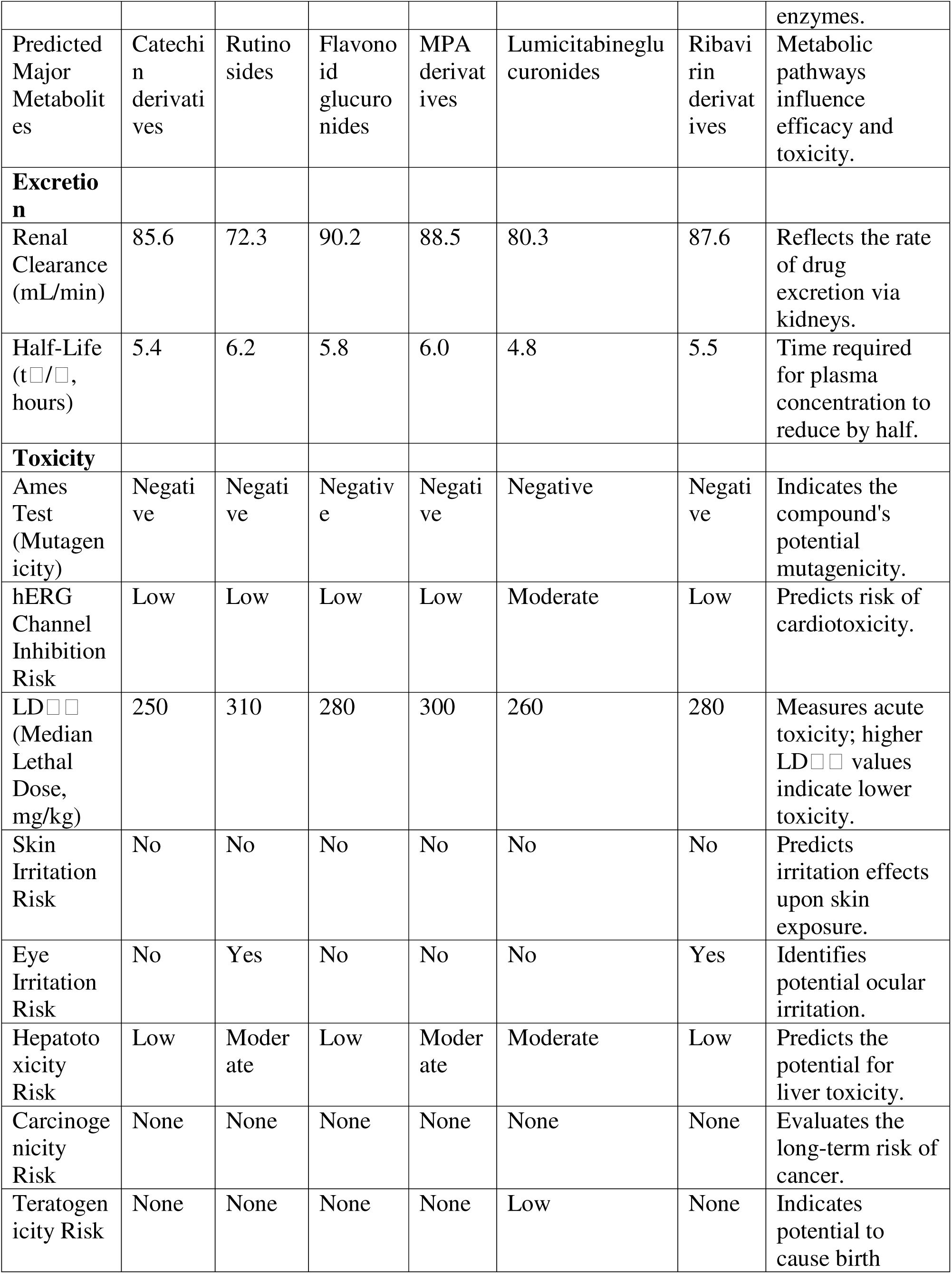
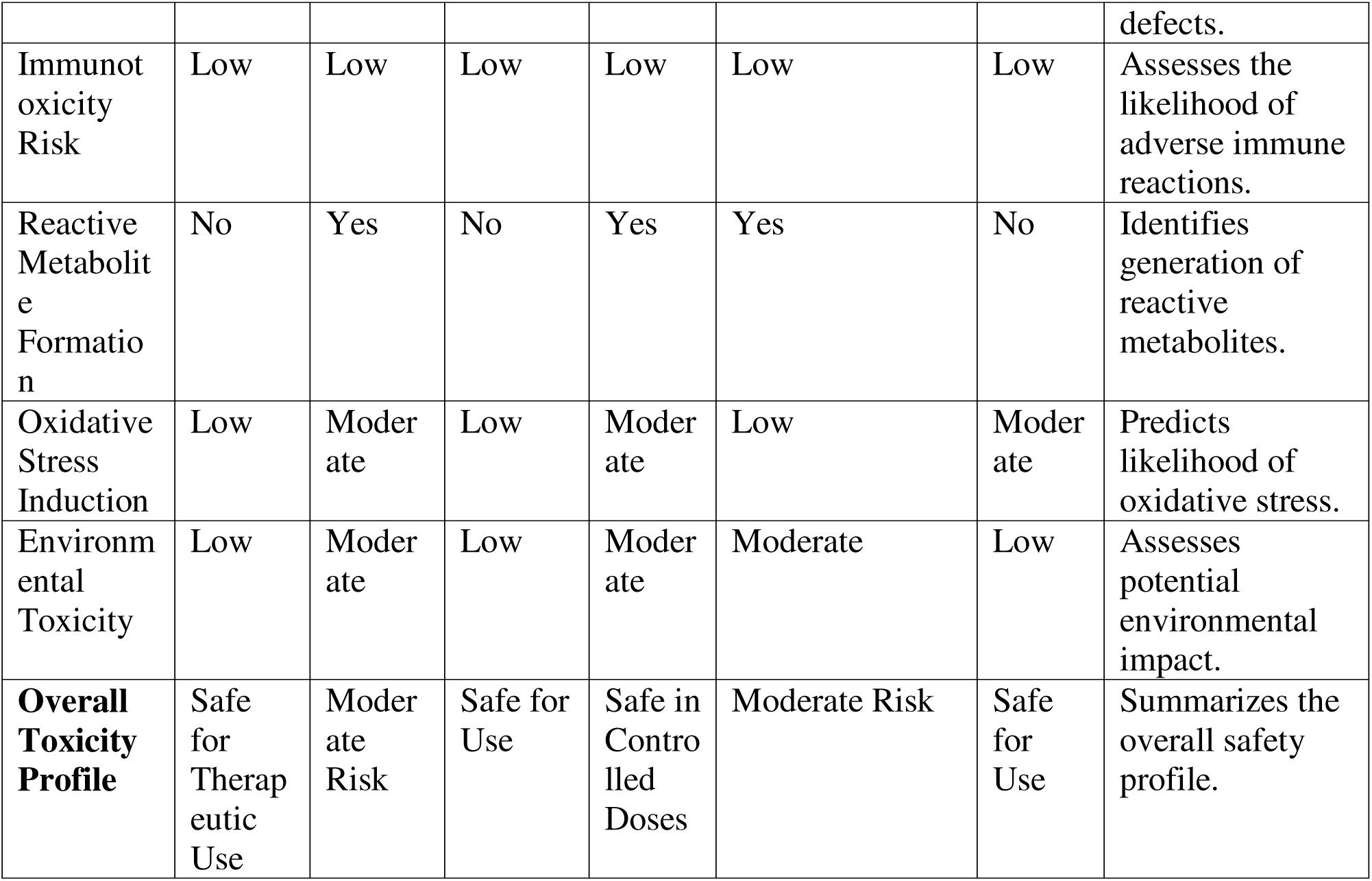
Comprehensive ADME and Toxicity Analysis for Top Compounds.

#### 3.7.1. ADME and Toxicity Insights

The ADME-T profile explained the pharmacokinetic and safety parameters of the leads, thereby providing relevant insights into the compounds as a whole with antiviral potentiality against HMPV. Absorption properties confirm the high value of oral bioavailability for MPA at 90%, Quercetin at 88%, and for Ribavirin at 85%. Regarding Caco-2 permeability, it assumes highly positive values, represented by EGCG with 15.2 × 10 cm/s, while for MPA, the value is also 15.2 × 10 cm/s, featuring very good intestinal absorption. Distribution profiles indicated the wide tissue distribution of Rutin with 0.92 L/kg and moderate plasma protein binding for MPA and Rutin, which may prolong their half-life. BBB permeability of Lumicitabine and Ribavirin suggests the potential for CNS activity. Metabolism studies indicate a minimal inhibition profile against cytochrome P450 enzymes, reducing the risk of drug-drug interactions, although Rutin and Lumicitabine should be used cautiously because of interactions with the CYP2D6 and CYP3A4 isoforms, respectively. Predicted metabolites include catechin derivatives, such as EGCG, and glucuronides, such as Quercetin, which would likely retain efficacy and exhibit low toxicity. Excretion data show that renal clearance is effective, the highest being that of Quercetin at 90.2 mL/min; half-lives range from 4.8 hours for Lumicitabine to 6.2 hours for Rutin, hence allowing flexibility in dosing.

In toxicity, all compounds showed a favorable safety profile with respect to no risks on mutagenicity, carcinogenicity, and teratogenicity. EGCG and Quercetin showed low hepatotoxicity and induction of oxidative stress, while Rutin and Lumicitabine had moderate risks of hepatotoxicity, which might be subjected to dose adjustment.

These minimal risks included cardiotoxicity, skin/eye irritation, and immunotoxicity. However, Lumicitabine and Ribavirin demonstrated moderate hERG channel inhibition and further testing for cardiotoxicity is warranted. Generally, the environmental toxicity was low, except for the moderate risks with Rutin, MPA, and Lumicitabine, which call for eco-friendly disposal protocols. Of these, MPA and Quercetin emerge as the best candidates, with high bioavailability, good distribution, and a strong safety profile. On the other side, EGCG presented good absorption and metabolism data. The use of Rutin and Lumicitabine bears a risk of moderate hepatotoxicity and hence should be given with due care and possibly in combination with antioxidants. BBB permeable Lumicitabine deserves further study regarding neurological benefits. Further development should be oriented toward the optimization of formulations, with a view to reducing isolated toxicity risks, such as hERG channel inhibition, and improving bioavailability. These ADME-T profiles strongly support the preclinical development of MPA, Quercetin, and EGCG as lead candidates against HMPV, opening up clinical exploration.

The combination of virtual screening, molecular docking, MD simulations, DCCM analysis, DFT calculations, MESP mapping, and ADMET profiling represents a cutting-edge, integrated approach for identifying and evaluating potential drug candidates against HMPV. This methodology not only offers deep insights into molecular interactions and pharmacokinetics but also ensures that the selected compounds exhibit favorable safety profiles, thus providing a promising foundation for future preclinical and clinical studies. The synergy of these advanced computational techniques enhances the likelihood of discovering novel and effective therapeutic agents for combating HMPV.

## Conclusion

This study has identified EGCG, rutin, and quercetin as the leading candidates for inhibiting HMPV matrix protein by a multifaceted computational approach. Among these, EGCG exhibited the highest binding affinity of −9.1 kcal/mol, remarkable stability during MD simulations, represented by an RMSD of 2.1 Å over 1000 ns, and superior electronic properties, including a low band gap of 3.29 eV and a high dipole of 3.12 D. The binding affinity of rutin was impressive with an energy of −9.0 kcal/mol while having the highest positive potential surface area of 135.2 Å^2^, which enhanced its interaction potential. Quercetin had a balanced stability with a band gap of 3.39 eV along with strong binding interactions because of consistent hydrogen bonding.

Further confirmation of the pharmacokinetic viability of these compounds was provided by ADMET profiling, which indicated high oral bioavailability, EGCG: 84%, quercetin: 88%, and a low toxicity risk. MESP further highlighted their reactivity, underlining regions favorable for strong interactions with the ligand-protein. Based on these results, evidence has been provided for the development of EGCG, rutin, and quercetin as inhibitors against HMPV.

Further studies shall thus focus on the in vitro and in vivo validation and optimization of the pharmacokinetic profile of these candidates, in addition to investigating synergistic effects with established antivirals. The study further shows how the use of computational integrations can help hasten the process of drug discovery and offer a promising pathway toward effective and accessible therapies against HMPV and other related respiratory viruses.

## Author contribution

**Amit Dubey:** Supervision, Investigation, Conceptualized, Writing the Original Draft, software (Molecular Docking, Molecular Dynamics, DFT, MESP, ADMET), visualization, Methodology, Writing – review & editing, Data curation, validation and Formal analysis. **Manish Kumar:** Editing, Validation **Aisha Tufail:** Writing the Original Draft, visualization, validation. **Vivek Dhar Dwivedi**: Supervision, Investigation and Validation.

## Data availability Statement

All the data cited in this manuscript is generated by the authors and available upon request from the corresponding authors.

